# Inhibiting a mRNA motif binding protein that mediates TGF-β1 upregulation of translation attenuates pulmonary fibrosis in mice

**DOI:** 10.1101/2022.10.23.513405

**Authors:** Wensheng Chen, Darrell Pilling, Richard H. Gomer

**Affiliations:** Department of Biology Texas A&M University, College Station, TX 77843-3474 USA

**Keywords:** TGF-β1, lung, NEU3, translation, mRNA, polysomes

## Abstract

In human lung cells, the profibrotic cytokine TGF-β1 increases sialidase 3 (NEU3) protein by increasing *NEU3* translation without increasing levels of *NEU3* mRNA. To elucidate how TGF-β1 regulates translation, we treated human lung fibroblasts (HLF) with TGF-β1 and used proteomics and RNA-seq to determine the effect of TGF-β1 on proteins, mRNAs, and mRNA polysome/monosome ratios. We identified 181 mRNAs where TGF-β1 also increases translation to increase protein levels without significantly affecting mRNA levels. These mRNAs share a common 20 nucleotide motif. Deletion or insertion of this motif in mRNAs eliminates or induces the TGF-β1 regulation of translation. At least 5 RNA-binding proteins including DDX3 bind the RNA motif, and TGF-β1 regulates their protein levels and/or binding to the motif. Inhibiting DDX3, either by siRNA or small molecule inhibitors, reduced TGF-β1 induced NEU3 levels. In the mouse bleomycin model of pulmonary fibrosis, injections of the DDX3 inhibitor RK-33 starting 10 days after bleomycin potentiated survival and reduced lung inflammation, fibrosis, and lung tissue levels of DDX3, TGF-β1, and NEU3. Together, these results suggest that TGF-β1 regulates RNA-binding proteins that interact with a mRNA motif that is necessary and sufficient for TGF-β1 to regulate mRNA translation, and that blocking this effect can reduce fibrosis.

## Introduction

Idiopathic pulmonary fibrosis (IPF) is a chronic and progressive lung disease with an average survival of 3–5 years following diagnosis (Abdul-Hafez et al., 2018; Cottin et al., 2021). Fibroblast and extracellular matrix accumulation replacing normal lung tissue in IPF patients leads to loss of lung function, disruption of gas exchange, and respiratory failure (Lederer and Martinez, 2018; Phan, 2002; Selman and Pardo, 2001). Sialic acid is often the distal sugar on glycoconjugates, and sialidase enzymes remove this sugar (Miyagi et al., 2012; Pshezhetsky and Ashmarina, 2013). The extracellular sialidase NEU3 is upregulated in fibrotic lesions in IPF patient lungs, and the lungs of mice with bleomycin-induced pulmonary fibrosis (Karhadkar et al., 2017). Mice lacking NEU3 essentially do not develop fibrosis in the bleomycin model, and aspiration of mouse NEU3 (but not enzyme-dead NEU3) causes pulmonary fibrosis in mice (Karhadkar et al., 2020a; Pilling et al., 2022). Inhibitors of NEU3 significantly reduce fibrosis in the mouse bleomycin model (Karhadkar et al., 2021; Karhadkar et al., 2017). Together, this suggests that NEU3 upregulation potentiates pulmonary fibrosis in mice.

The cytokine transforming growth factor-β1 (TGF-β1) is sequestered in an inactive state in the extracellular environment by being effectively wrapped in 2 copies of a protein called LAP (Hinck et al., 2016; Robertson and Rifkin, 2016), and high levels of the released active form of TGF-β1 are often observed in tissues during inflammation, and in diseases such as cancer and fibrosis (Akhurst and Hata, 2012; Derynck and Akhurst, 2007). The high extracellular levels of active TGF-β1 drive disease progression by modulating cell growth, migration, or changes in cell morphology and/or phenotype (Derynck and Akhurst, 2007; Freimuth et al., 2012). High levels of extracellular TGF-β1 are present in the fibrotic lesions in IPF, and the TGF-β1 activates receptors on fibroblasts to cause them to proliferate and increase the production of extracellular matrix, as well as the production of NEU3 (Karhadkar et al., 2017; Phan, 2002; Smith et al., 2015).

The TGF-β receptor activates a signal transduction pathway that regulates SMAD proteins, which in turn can regulate the transcription of specific genes (Davis-Dusenbery and Hata, 2011; Shi and Massagué, 2003). In addition, TGF-βs can regulate translation through a variety of mechanisms. In human pancreatic cancer cells, SMAD4 can promote or inhibit mRNA entry into polysomes, although the mechanism is unknown (Thornley et al., 2012). TGF-β1 regulates epithelial–mesenchymal transition (EMT) in part by phosphorylation of the translation initiation factor eIF4F (Smith et al., 2015). In human breast epithelial cells, the TGF-β1-SMAD pathway decreases translation of the p53 mRNA by inhibiting the binding of the p53 mRNA to a ribosomal protein and a translation elongation factor (López-Díaz et al., 2013). In human cardiac fibroblasts incubated with TGF-β1 and in human dilated cardiomyopathy tissue, RNA-seq of monosomes, polysomes, and total mRNA indicated that one-third of the proteins regulated by TGF-β1 are regulated at the level of translation (Chothani et al., 2019). We previously found that TGF-β1 increases NEU3 protein levels and increases the amount of *NEU3* mRNA in polysomes without affecting *NEU3* total mRNA levels in human lung epithelial cells (Chen et al., 2020). This indicated that TGF-β1 increases levels of NEU3 at the level of translation and not transcription.

RNA binding proteins (RBPs) are capable of regulating translation for the rapid modulation of protein synthesis from the existing mRNA (Hentze et al., 2018). RBPs can either sequester mRNA to slow translation, such as the P-body proteins AGO2 and UPF1; promote translation, such as DDX3, or actively suppress and degrade mRNA, such as CELF2 (Hubstenberger et al., 2017; Mo et al., 2021; Schneider-Lunitz et al., 2021). DDX3 overexpression has been implicated in many diseases including viral infections, inflammatory autoimmune diseases, and many forms of cancer (Bol et al., 2015; Kukhanova et al., 2020; Tantravedi et al., 2019).

In this report, we identified proteins in addition to NEU3 in human lung fibroblasts whose levels are increased by TGF-β1 at the level of translation but not transcription, and identified a common motif in the encoding mRNAs. We then found that the motif is necessary and sufficient for the TGF-β1 regulation of translation, and identified RNA-binding proteins that bind to, or decrease binding to, the motif in response to TGF-β1. A protein that binds the mRNA motif in response to TGF-β1 is DDX3, and downregulation of DDX3 or treatment with the DDX3 inhibitor RK-33 reduced TGF-β1 induced NEU3 levels in fibroblasts. In a mouse model of pulmonary fibrosis, RK-33 reduced bleomycin-induced lung inflammation and fibrosis, suggesting that the TGF-β1 upregulation of translation pathway may be a viable target to inhibit fibrosis.

## Materials and Methods

### Cell culture and TGF-β1 treatment

Four human lung fibroblast (HLF) cell lines, from 2 different healthy males, one healthy female, and one female IPF patient, were gifts from Dr. Carol Feghali-Bostwick, Medical University of South Carolina. Cells were cultured in DMEM (#12-614F, Lonza, Walkersville, MD) supplemented with 10% bovine calf serum (BCS) (Seradigm, Randor, PA), 100 U/ml penicillin, 100 μg/ml streptomycin, and 2 mM glutamine (all from Lonza). For all experiments, cells were used at passages 4–8. 5 × 10^5^ HLF were cultured in 100 mm tissue culture dishes (Corning, Corning, NY) for 24-48 hours. When the cell density reached ∼60% confluence, the supernatant was removed and cells were gently rinsed with prewarmed PBS 3 times. The medium was changed to prewarmed 10 ml DMEM supplemented with 100 U/ml penicillin, 100 μg/ml streptomycin, and 2 mM glutamine (no BCS) with or without 10 ng/ml TGF-β1 (PeproTech, Rocky Hill, NJ). After 48 hours, the cell density reached 80–90% confluence, and cells were rinsed with 10 ml PBS three times before detaching with 1 ml Accutase (Innovative Cell Technologies, San Diego, CA) for 5 minutes. Cells were collected, counted, and equal number of cells were lysed for the following assays.

### Proteomics

2 x 10^6^ HLF were collected by centrifugation at 300 x g for 5 minutes at 4°C. The supernatant was discarded and cells were washed twice with 1 ml ice-cold PBS. The cell pellet was resuspended in 0.3 ml of ice-cold RIPA Buffer (Cat# 89900, ThermoScientific, Rockford, IL) with 1x Protease and Phosphatase Inhibitor (PPI) Cocktail (Cat# 78440, ThermoScientific, Rockford, IL). Aliquots of the lysate were then snap frozen in liquid nitrogen and stored at −80°C. In-gel protein preparation was performed as described on the University of Texas-Southwestern Proteomics Core webpage (https://proteomics.swmed.edu/wordpress/?page_id=553). Briefly, 50 µg of protein per test sample was combined with 5x SDS sample buffer (250 mM Tris-HCl, pH 6.8, 10% SDS, 30% glycerol, 0.02% bromophenol blue dye, and 5% β-mercaptoethanol) and heated at 95°C for 10 minutes. 40 µg of protein was loaded onto a 4-20% Tris/glycine gel (Bio-Rad, Hercules, CA, USA), and run at 70V for ∼20 minutes (until ∼5-10 mm through the gel). The gel was stained with fresh Coomassie blue dye (Cat# 24615, ThermoScientific, Rockford, IL) for 1 hour with gentle shaking, destained with buffer (10% acetic acid, 40% methanol, and 50% ddH_2_O) for 1.5 hours, and further destained overnight in sterile water. In a glass petri dish, the protein bands were cut into 1-2mm cubes, and transferred to 1.5 mL microcentrifuge tubes. Samples were sent on ice to the University of Texas Southwestern Proteomics Core (https://proteomics.swmed.edu/wordpress/) for trypsin digestion and Thermo Fusion Lumos standard gradient mass spectrometry. At the Core, individual peptides were assigned to proteins using Proteome Discoverer 2.4 (Orsburn, 2021) using the human protein database from UniProt (The UniProt, 2015; The UniProt, 2021), and Proteome Discoverer then generated abundances of proteins in each sample.

### Ribosomal RNA collection, purification, and sequencing

10-20 x 10^6^ HLFs were collected and treated as described previously for monosome and polysome fractionation (Chen et al., 2020). Lysed samples were separated on a 10-50% sucrose gradient made with “Polysome Gradient Buffer” (10 mM HEPES-KOH pH 7.5, 70 mM NH_4_OAc, 5 mM Mg(OAc)_2_) and the associated (w/v) sucrose solutions prepared the same day. Cell lysates were layered on top of the prepared sucrose gradient and centrifuged and fractionated following the manufacturer’s instructions for a TRiAX flow cell (BioComp Instruments, New Brunswick, Canada) and FC203B fraction collector (Gilson, Middleton, WI). RNA purification and precipitation was performed as described (Chen et al., 2020). In addition, total RNA was purified as previously described (Chen et al., 2020). RNA pellets were air dried for at least 10 minutes at room temperature until pellets were completely dry before being dissolved in 20 µl nuclease-free water (ThermoScientific, Rockford, IL). RNA concentration was checked using a Synergy Mx plate reader with a Take3 microvolume plate (BioTek, Winooski, VT). RNA-seq libraries were created following the manufacturer’s instructions using QuantSeq 3’ mRNA-Seq FWD library prep kits for Illumina (Lexogen Inc, Greenland, NH), with 2 µg of RNA used as the starting material. Libraries were sequenced using an Illumina NextSeq 500 platform (Texas A&M University Institute for Genome Sciences and Society, College Station, TX). RNA sequencing data was analyzed using the QuantSeq Data Analysis Pipeline on the BlueBee Genomic Platform (BlueBee, San Mateo, CA).

### Proteomics and genomics data analysis

Protein or RNA reads from the control and TGF-β1 group (n=4) were compared by t-test. For the significantly changed proteins or RNAs (p < 0.05), the TGF-β1 / control ratios of the mean reads were calculated. Proteins or RNAs with a ratio > 1 were defined as “Up” (statistically increased in the TGF-β1 group), and proteins or RNAs with a ratio < 1 were defined as “Down” (statistically decreased in TGF-β1 group). For the proteins or RNAs that were not significantly changed, the 95% confidence interval (95% CI) of the reads were compared between control and TGF-β1 group (Bristol, 1999; Stegner et al., 1996; Wellek, 2002). Briefly, if the 95% CI of a protein or RNA in TGF-β1 group was completely within the range defined as plus and minus 30% of the reads in the control group, the reads of this protein or RNA were considered statistically equivalent, and this protein or RNA was defined as “Stable”. The non-changed and non-stable proteins or RNAs were defined as “Others”.

For the ribosomal mRNAs (n=4), a Polysome / Monosome ratio (PM ratio) was calculated and compared between control and TGF-β1 group with t-test. For the PM ratio significantly changed mRNAs (p < 0.05), an X value was calculated. mRNAs with X > 1 were defined as “M to P shifted” (PM ratio significantly increased in TGF-β1 group), and mRNAs with X < 1 were defined as “P to M shifted” (PM ratio significantly decreased in TGF-β1 group). For the mRNAs that did not show a PM ratio change, the 95% confidence interval (95% CI) of the PM ratios were compared. If the 95% CI of a mRNA PM ratio in TGF-β1 group was completely within the range defined as plus and minus 30% of the PM ratio in the control group, the PM ratio of this mRNA was considered statistically equivalent, and this protein was defined as “M/P Stable”. The non-shifted and non-stable mRNAs were defined as “Others”.

### GO analysis

Gene ontology (GO) biological process analysis was performed using GO Panther (v17.0) (Ashburner et al., 2000; The Gene Ontology, 2021) and analysis was confirmed and graphs generated by ShinyGO (v 0.74 using Ensembl 92 Release 104) (Ge et al., 2020). Groups were analyzed compared to the standard ‘all proteins in the *Homo sapiens* database’, and significance (p < 0.05) was determined by Fisher’s exact test with False Discovery Rate (FDR) correction.

### Common motif and uORF analysis

mRNA sequences with 5’ untranslated region (5’UTR), coding sequence (CDS) and 3’ untranslated region (3’UTR) were downloaded from the UCSC Genome Browser (http://genome.ucsc.edu). Common motifs of the mRNAs were identified using MEME (Bailey et al., 2015). Briefly, mRNA sequences were uploaded to MEME as input, and common motifs were searched for using the ‘zoops’ parameter and 10 to 25 nucleotide motifs. Upstream open reading frame (uORF) analysis was performed at NCBI ORFfinder (https://www.ncbi.nlm.nih.gov/orffinder/) with “ATG only” and “ATG and alternative initiation codons” settings.

### Plasmid construction, transfection and qPCR

The human Myc-NEU3 expression plasmid RC216537 (Origene, Rockville, MD) was used to express NEU3. Myc-NEU3 mutants were generated using a QuikChange II Site-Directed Mutagenesis Kit (Agilent, Santa Clara, CA) with the DNA oligomers GGGCCCCTTAAACCACTTATTGAATCCACACTACC for mutant 1 and CAGTTCACTTAGACTGGAAGATGAATCTGGAACAC for mutant 2, (supplemental file 7), and the resulting plasmids were sequenced to confirm the point mutations and absence of other mutations. Human Myc-ESD and Myc-MKKS expression plasmids were constructed at VectorBuilder (Chicago, IL) (supplemental files 8 and 9).

1 × 10^5^ HLFs were mixed with 2 μg of 100 μg/ml of the above expression plasmids in 100 μl PBS (GE Lifesciences, Marlborough, MA) and were transfected by electroporation using a 4D-Nucleofector System (Lonza, Walkersville, MD) following the manufacturer’s protocol CZ-167. The transfected cells were kept at room temperature for 15 minutes for recovery, after which the cells were cultured in 10 ml DMEM with 10% BCS and 2 mM glutamine in a humidified incubator at 37°C with 5% CO_2_. After 24 hours, 400 μg/ml of G418 (Calbiochem EMD, San Diego, CA) was added to select for transfected cells. The culture medium with G418 was refreshed every 3 or 4 days. After 14 days, the cells were collected and used for assays described below. RNA extraction from the transfected cells and qPCR for Myc-NEU3, Myc-ESD and Myc-MKKS was done as we described previously (Chen et al., 2020) using the primers described in supplemental files 7, 8 and 9.

### RNA-binding protein prediction and RNA-protein interaction assay

RNA-binding proteins (RBPs) that can interact with Group 4 mRNAs were identified using CLIP-seq databases on POSTAR3 (http://111.198.139.65/RBS.html) with a RBP Binding Sites Module. RNA names of Group 4 mRNAs were entered and interacting RBPs were discovered from HIT-CLIP and iCLIP databases (Hu et al., 2017; Yang et al., 2015; Zhao et al., 2022; Zhu et al., 2019). The interaction probabilities of the Group 4 motif GGAGGAGGAGGAGGAGGAGG and RNA-binding proteins was predicted using RPIseq (http://pridb.gdcb.iastate.edu/RPISeq/<u>#</u>). For this analysis, the sequence of the motif and the sequence of each possible RBP was entered, and the interaction probability was predicted by a Random Forest classifier and a Support Vector Machine classifier (Muppirala et al., 2011; Muppirala et al., 2013).

Biotin-labeled Group 4 motif (Bio-motif; supplemental file 10) was synthesized at IDT (Coralville, IA). 5 μl of a 50% slurry of streptavidin agarose beads (Millipore, Billerica, MA) was gently mixed with 500 μl ice-cold streptavidin agarose wash buffer (SWB) (20 mM Tris-HCl pH 7.5, 100 mM KCl, 2 mM EDTA, 0.5 mM DTT, 0.5 mM PMSF) in a 1.5 ml Eppendorf tube. The beads were collected by centrifugation at 500 x g for 2 minutes at 4℃, resuspended in 500 μl ice-cold SWB, and 1 μl (1000 pmol) of Bio-Motif was added to the tube and incubated with the beads at 4℃ for 2 hours with 100 RPM rotation. The beads were collected by centrifugation at 500 x g for 2 minutes at 4℃. Beads were washed three times by resuspension in 500 μl Bio-Motif-Streptavidin interaction buffer (SIB) (20 mM Tris-HCl pH 7.5, 300 mM KCl, 0.2 mM EDTA, 0.5 mM DTT, 0.5 mM PMSF) followed by centrifugation. Beads were then resuspended in 500 μl SIB after the last rinse. Approximately 1 × 10^6^ cells treated with or without TGF-β1 as above were lysed with 200 µl RIPA Buffer with 1x Protease and Phosphatase Inhibitor Cocktail at 4℃ for 30 minutes, and the mixture was clarified by centrifugation at 12,000 x g for 10 minutes at 4℃. 100 µl of the supernatant (protein concentration at 3 μg/μl) was added to the beads and incubated at 4℃ overnight with 100 RPM rotation. The beads were then collected by centrifugation at 500 x g for 2 minutes at 4℃ and washed three times as described above with 400 μl SIB. Beads were resuspended in 40 μl SIB, and 10 μl 5x SDS sample buffer was added and the mixture was heated at 95℃ for 10 minutes. After cooling, this was clarified by centrifugation at 10,000 x g for 10 minutes at room temperature. The supernatant was collected and used for Coomassie blue staining and western blotting.

### DDX3 siRNA transfection and DDX3 inhibitor treatment in HLF

HLFs were transfected with DDX3 siRNA (#4392420, Thermo Scientific, Rockford, IL) or negative control siRNA (#4390843, Thermo Scientific), following the manufacturer’s instructions. After 24 hours, the transfection medium was removed and HLFs were treated with or without 10 ng/ml TGF-β1 in no serum added DMEM. After an additional 48 hours, the culture medium was discarded and cells were lysed in RIPA buffer with PPI cocktail. Samples were analyzed by Coomassie blue staining and western blotting. Additionally, HLFs were treated with 1 or 10 μM RK-33 (Selleck Chemicals, Houston, TX), or 1 or 10 μM IN-1 (AdooQ, Irvine, CA) from a 10 mM stock in DMSO (VWR) or DMSO diluent control, in the presence or absence of 10 ng/ml TGF-β1 (PeproTech, Rocky Hill, NJ). After an additional 48 hours, the culture medium was discarded and cells were lysed in RIPA buffer with PPI cocktail. Samples were analyzed by Coomassie blue staining and western blotting.

## Pulmonary fibrosis mouse model and RK-33 treatment

This study was carried out in strict accordance with the recommendations in the Guide for the Care and Use of Laboratory Animals of the National Institutes of Health. The protocol was approved by Texas A&M University Animal Use and Care Committee (IACUC 2020-0272). All procedures were performed under anesthesia, and all efforts were made to minimize suffering. To induce inflammation and fibrosis, 7-8 week old 20-25 g male C57BL/6 mice (Jackson Laboratories, Bar Harbor, ME) were given an oropharyngeal aspiration of 4.5 U/kg bleomycin (Cat# BML-AP302-0010, Enzo Life Sciences, Farmingdale, NY) in 50 µl of 0.9% saline or oropharyngeal saline alone, as a control, as previously described (Karhadkar et al., 2020a; Pilling et al., 2022). All the mice were monitored twice daily to observe any sign of distress. At days 3, 7, 10, 14 and 21 after bleomycin treatment, mice were euthanized by CO_2_ inhalation. Bronchoalveolar lavage fluid (BALF) was obtained and BALF cells counted, as described previously (Karhadkar et al., 2020a; Pilling et al., 2022). After collecting BALF, the lungs were inflated with, and embedded in, OCT compound (#3801480, Leica, Buffalo Grove, IL), frozen, and stored at −80 °C. 10 µm cryosections were cut on a CM1520 cryostat (Leica Biosystems, Deer Park, IL) and used for staining as described below.

For mice receiving RK-33 treatment, inflammation and fibrosis were induced as described above. At days 10, 12, 14, 16, 18 and 20 after saline or bleomycin treatment, mice were given daily intraperitoneal injections of 20 mg/kg RK-33 (formulated as 40 mg/ml in DMSO) diluted in saline, or 100 µl DMSO/saline diluent control. Mice were euthanized at Day 21 after bleomycin treatment. Bronchoalveolar lavage fluid (BALF) and lung tissue was obtained and analyzed as described above.

### Bronchoalveolar lavage fluid (BALF) cell counting

The BALF cells were clarified and collected by centrifugation at 500 x g for 10 minutes, and total BALF cell numbers were counted. BALF cell spots were prepared and stained with Wright–Giemsa stain (#08711, Polysciences, Inc., Warrington, PA) as previously described (Karhadkar et al., 2020a). Using a 40x objective, at least 150 cells from each stained BALF spot were examined and the percent positive cells was recorded.

### PicroSirius red stain and hydroxyproline assay

Lung sections were stained with Sirius red to detect collagen and analyzed as previously described (Karhadkar et al., 2020a). Hydroxyproline assays were performed as described previously (Karhadkar et al., 2020a; Karhadkar et al., 2021). Briefly, approximately half of lobes of lungs frozen in OCT were cut off in three pieces, thawed, and washed three times with 1 ml PBS to remove OCT by centrifugation at 2000g for 5 minutes in preweighed Eppendorf tubes. After the last centrifugation, the tubes were kept inverted for 5 minutes to allow PBS to blot onto blotting paper, and the tissue was then weighed. Tissues were then processed using a hydroxyproline quantification kit (MAK008-1KT, Sigma-Aldrich) following the manufacturer’s directions.

### Immunofluorescence and immunohistochemistry

The Myc-NEU3, Myc-ESD, and Myc-MKKS transfected and G418 selected HLFs were seeded on 96 well black μ-Plates (Cat# 89626, ibidi USA, Fitchburg, WI) at 2000 cells per well. Cells were treated with or without 10 ng/ml TGF-β1 for 48 hours at 37℃ in a humidified 5% CO_2_ incubator. Cells were fixed with 4% paraformaldehyde (PFA) in PBS for 10 minutes as previously described (Rijal et al., 2020). Cells were rinsed with PBS three times and permeabilized with 0.5% Triton-X-100 in PBS for 5 minutes. After three PBS rinses, cells were blocked with 2% BSA in PBS (PBSB) for 1 hour at room temperature. 5 μg/ml rabbit anti-Myc-Tag antibody (Cat# 2278, Cell Signaling Technology, Danvers, MA) in PBSB was incubated with cells overnight at 4 ℃. Cells were washed three times for 5 minutes per wash with PBS and stained with 1 μg/ml DAPI dye (BioLegend, San Diego, CA) and 2 μg/ml donkey F(ab)_2_ anti-rabbit Alexa Fluor 488 (Jackson ImmunoResearch, West Grove, PA) or 5 μg/ml donkey F(ab)_2_ anti-rabbit Rhodamine Red-X antibody (Jackson ImmunoResearch) in PBSB at room temperature for 30 minutes. Cells were washed three times with PBS, and images were taken with a Nikon ECLIPSE Ti2 microscope (Nikon, Melville, NY).

Cryosections of mouse lungs were fixed in acetone for 20 minutes at room temperature and then rehydrated in water for 5 minutes and then PBS for 5 minutes. Slides were blocked with PBSB for 30 minutes and stained with 5 μg/ml rabbit antibodies in PBSB at 4℃ overnight. Rabbit antibodies were either anti-TGF-β1 (NBP1-45891), anti-GSTO1 (NBP2-32691), anti-IGBP1 (NBP1-83126), anti-MAGI3 (NBP1-81266), anti-MTCL1 (NBP2-47366), anti-LHFPL2 (NBP1-93654), anti-RBM33 (NBP1-84200), anti-BAF180 (NB100-79833), anti-APC4 (NBP1-90137) (all from Novus Biologicals, Centennial, CO), and anti-NEU3 (27879-1-AP, Proteintech, Rosemont, IL). Monoclonal antibodies at 5 μg/ml were rat anti-CD45 (Cat#304002, BioLegend), rat anti-CD11b (Cat#101202, BioLegend), and hamster anti-CD11c (Cat#117301, BioLegend). Directly conjugated antibodies at 5 μg/ml were Alexa Fluor 647 conjugated anti-CD45 (Cat#103123, BioLegend), Alexa Fluor 488 conjugated anti-TGF-β1 (NBP2-45137AF488, NOVUS) and Alexa Fluor 647 conjugated anti-DDX3 (NBP2-14848AF647, NOVUS). The staining was further processed as described previously (Cox et al., 2015; Karhadkar et al., 2019). Briefly, slides were incubated with combinations of 2 μg/ml donkey F(ab)_2_ anti-rabbit Alexa Fluor 488 (Cat# 711-546-152, Jackson ImmunoResearch), 2 μg/ml goat F(ab)_2_ anti-hamster Alexa Fluor 488 (Cat# 107-546-142, Jackson ImmunoResearch), 5 μg/ml donkey F(ab)_2_ anti-rat Rhodamine Red-X antibody (Cat# 712-296-153Jackson ImmunoResearch) and 5 μg/ml donkey F(ab)_2_ anti-rabbit Rhodamine Red-X antibody (Cat# 711-296-152, Jackson ImmunoResearch) in PBSB at room temperature for 30 minutes. The staining was further processed as described previously (Cox et al., 2015; Karhadkar et al., 2019). The slides were kept in the dark at 4°C for 1 h to harden the antifade mounting media with DAPI (Cat# H-1500, Vector).

Human lung tissue sections were obtained from the National Heart Lung and Blood Institute-sponsored Lung Tissue Research Consortium (LTRC). Formalin-fixed paraffin-embedded slides were dewaxed with xylene, then rehydrated through a graded series of alcohols, distilled water, and then PBS (Pilling et al., 2009). Slides were then blocked by incubation in PBSB for 60 minutes. Endogenous biotin was blocked by the addition of streptavidin and biotin solutions, following the manufacturers’ instructions (Streptavidin/Biotin Blocking Kit, Vector Laboratories, Burlingame, CA). Slides were then incubated overnight at 4°C with 1 μg/ml rabbit anti-DDX3 (NBP1-85291, Novus) in PBSB. Primary antibodies were detected with 1 µg/ml biotinylated donkey F(ab’)2 anti-rabbit IgG (711-066-152; Jackson ImmunoResearch) in PBSB for 30 minutes. Biotinylated antibodies were detected by a 1/500 dilution of ExtrAvidin alkaline phosphatase (Vector Laboratories) in PBSB for 30 minutes. Staining was developed with the Vector Red Alkaline Phosphatase Kit (Vector Laboratories) for 10 minutes. Sections were then counterstained for 30 seconds with Gill’s hematoxylin #3 (Sigma-Aldrich). Slides were mounted with VectaMount (Vector Laboratories). Tissue sections stained with antibodies were imaged with a Nikon Eclipse Ti2 microscope (Nikon Instruments, Melville, NY) and analyzed with ImageJ2 software (Rueden et al., 2017). The percentage area of tissue stained with was quantified as a percentage of the total area of the tissue, as described previously (Karhadkar et al., 2020b; Karhadkar et al., 2017).

### Coomassie blue staining and western blotting

Approximately 10^6^ cells were collected by centrifugation as described above and lysed in 200 μl RIPA Buffer (ThermoScientific) with 1x Protease and Phosphatase Inhibitor Cocktail (ThermoScientific). 80 μl of cell lysate was mixed with 20 μl 5x SDS protein loading buffer and heated at 95℃ for 5 minutes. Equal volumes of the heated samples were loaded on 4–20% Tris– glycine Mini-Protean TGX gels (Bio-Rad, Hercules, CA) for SDS-PAGE. The gel was stained with freshly prepared Coomassie blue dye for 1 hour with gentle shaking and destained overnight. Western blots were done as previously described (Chen et al., 2020), staining with 0.5 μg/ml ribbit anti-Myc-Tag antibody (Cell Signaling Technology). Where indicated, blots were stained with 1 μg/ml rabbit antibodies to AGO2 (NBP2-67121), UPF1 (NBP1-89641), DDX3 (NBP1-85291), U2AF2 (NBP2-58989), CELF2 (NBP2-16035), ALYREF (NBP1-90179), GSTO1 (NBP2-32691), IGBP1 (NBP1-83126), MAGI3 (NBP1-81266), LHFPL1 (NBP2-47366), RBM33 (NBP1-84200), BAF180 (NB100-79833) (all from Novus Biologicals), p-AGO2 (AP5291, ECM Biosciences, Versailles, KY), p-UPF1 (07-1016, Millipore, Billerica, MA) or NEU3 (27879-1-AP, Proteintech). Peroxidase-conjugated donkey F(ab)_2_ anti-rabbit (Cat# 711-036-152, Jackson ImmunoResearch) was used as the secondary antibody. SuperSignal West Pico Chemiluminescence Substrate (Thermo Scientific, Rockford, IL) was used following the manufacturer’s protocol to visualize the peroxidase using a ChemiDoc XRS+ System (Bio-Rad).

### Statistical Analysis

Statistical analysis was performed using Prism v7 (GraphPad Software, La Jolla, CA). Statistical significance between two groups was determined by t test, or between multiple groups using analysis of variance (ANOVA) with Dunnett’s post-test, and significance was defined as p<0.05. For the multiple t tests in the proteomics and RNA-seq data analysis, a false discovery rate (FDR) approach was used with Q=0.05. Statistical equivalence between two groups was determined by the 95% confidence intervals.

## Results

### TGF-β1 induces ribosomal mRNA shifts and leads to mRNA/protein mismatches in human lung fibroblasts

To elucidate the effect of TGF-β1 on fibroblasts, four human lung fibroblast cell lines were treated with or without recombinant human TGF-β1. Proteomics and RNA-seq data were analyzed according to the workflow in Figures 1 A and B. Proteomics identified 3,309 proteins, and TGF-β1 increased levels of 1,044 of these proteins and decreased levels of 44 proteins (p<0.05; multiple 2-tail t tests, n=4), and 714 proteins were statistically equivalent with or without TGF-β1 treatment (Figure 1c). In agreement with work from other labs (Mullenbrock et al., 2018; Senavirathna et al., 2020; Souma et al., 2018), RNA-seq analysis of these cells identified 22,466 different transcripts (a mix of mRNAs and non-coding RNAs), and TGF-β1 increased levels of 1,436 RNAs and decreased levels of 1,567 RNAs (p<0.05; multiple 2-tail t tests, n=4), while 3,556 RNAs were statistically equivalent with or without TGF-β1 treatment (Figure 1c). As we did for lung epithelial cells (Chen et al., 2020), control and TGF-β1-treated fibroblasts were fractionated to obtain free mRNA and monosome-associated mRNA (henceforth referred to as monosome fraction), and polysome-associated mRNA. As with the analysis of the proteomics and RNA-seq of total mRNA, multiple t tests were used to identify mRNAs with a statistically significant TGF-β1-induced shift from monosomes to polysomes, a shift from polysomes to monosomes, or a statistically significant absence of a shift (Figure 1b and c). Some protein/mRNA pairs did not show statistical significance for increase, decrease, or no change in one or more of the categories (mRNA, polysome/monosome ratio, or protein) and were thus excluded from further analysis. Combining the data, there were 1,149 proteins with statistical significance (increase, decrease, or no change) for the TGF-β1 effect on mRNA, polysome/monosome ratio, and protein.

**Figure 1.**
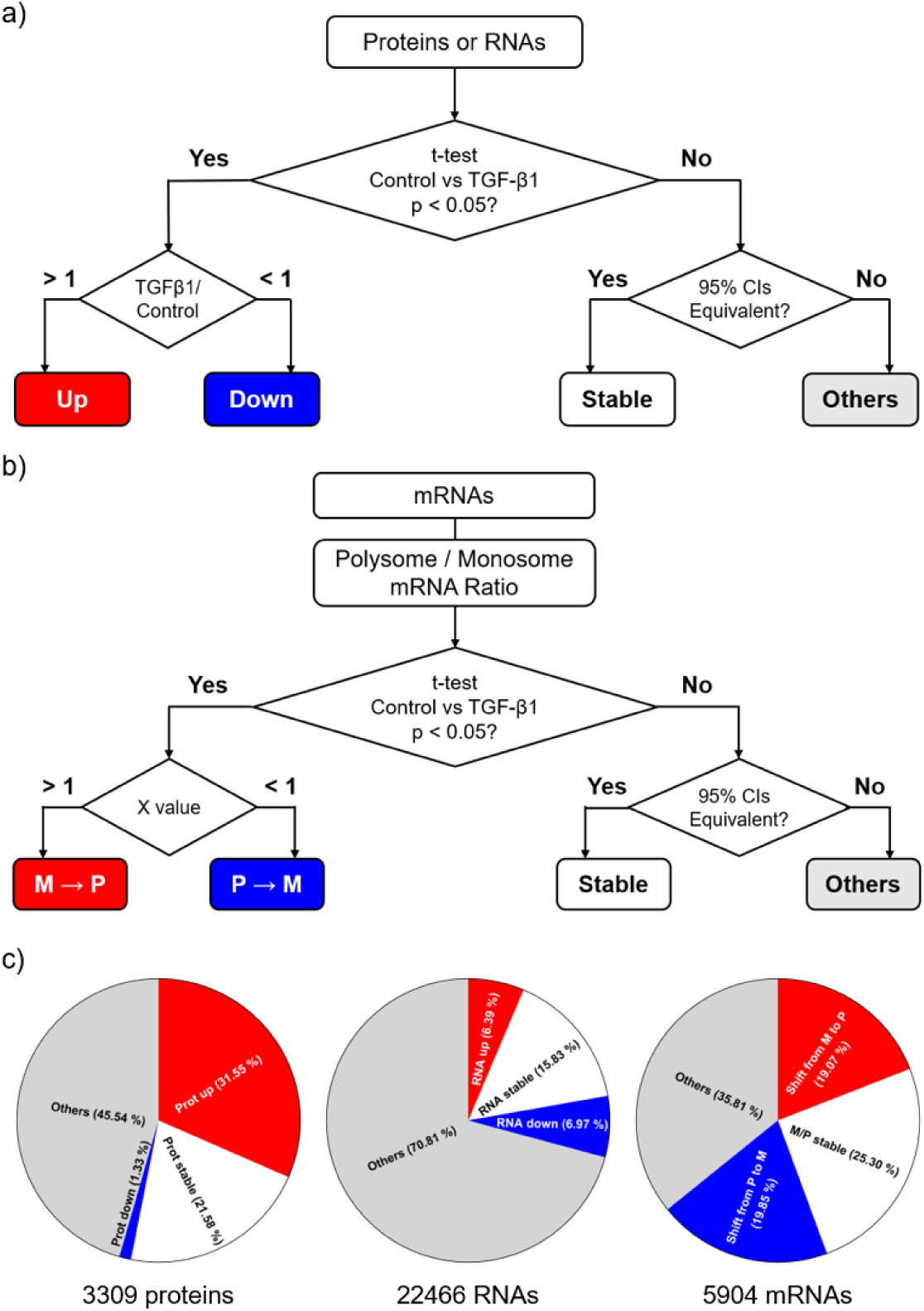
Overview of the data. **a-b)** Strategy of proteomics and RNA-seq analysis of 4 HLF cell lines treated with or without TGF-β1, starting with t-tests to compare the effect of TGF-β1 to control for each protein, RNA, or polysome/monosome ratio for a mRNA. In **b**, MèP indicates a TGF-β1-induced shift from monosomes to polysomes (and thus a TGF-β1-induced increased in translation) for a mRNA; PèM indicates a TGF-β1-induced shift from polysomes to monosomes. **c)** Distribution of protein responses to TGF-β1, distribution of RNA responses to TGF-β1, and distribution of polysome/monosome ratios for mRNAs detected in monosomes and/or polysomes.

The 1,149 proteins were divided into 27 groups according to total protein levels, total mRNA levels, and ribosomal mRNA shifts (Table 1 and supplementary files 1, 2, 3, 4, 5 and 6). TGF-β1 increased levels of 693 and decreased levels of 27 of the 1,149 proteins. For 545 of the 1,149 proteins, the TGF-β1 effect on the protein level did not match the TGF-β1 effect on the mRNA level (groups 4-12 and 16-24). For the proteins upregulated by TGF-β1, GO analysis indicated that Group 1 includes extracellular matrix and amino acid biosynthesis. Group 2 angiogenic factors and exocytosis, and Group 3 protein transport and regulation of ATPase activity (Figure S1). Group 4 (total mRNA stable • translation up • protein up) proteins include 26S protease regulatory subunits, ribosome proteins, and translation initiation and elongation factors. Groups 5 and 6 include signal recognition particle and nonsense-mediated decay proteins. Group 7 (total mRNA down • translation up • protein up) includes cellular stress response and autophagy proteins. Groups 19-27 (proteins decreased by TGF-β1) all had less than 10 proteins, and did not show any significant enrichment. Although NEU3 had no proteomics reads, it is a group 4-like protein (TGF-β1 increases NEU3 protein in lung fibroblasts (Karhadkar et al., 2017)), there was no significant effect of TGF-β1 on total mRNA level (control 26.7 ± 4.5, TGF-β1 24.9 ± 3.6, both in units of relative abundance from the RNA-seq, n=4) and *NEU3* showed a monosome to polysome mRNA shift (X = 5.92, p < 0.01). Together, these results suggest that TGF-β1 has a wide and complex effect on both transcription and translation control of human lung fibroblasts.

**Table 1.**
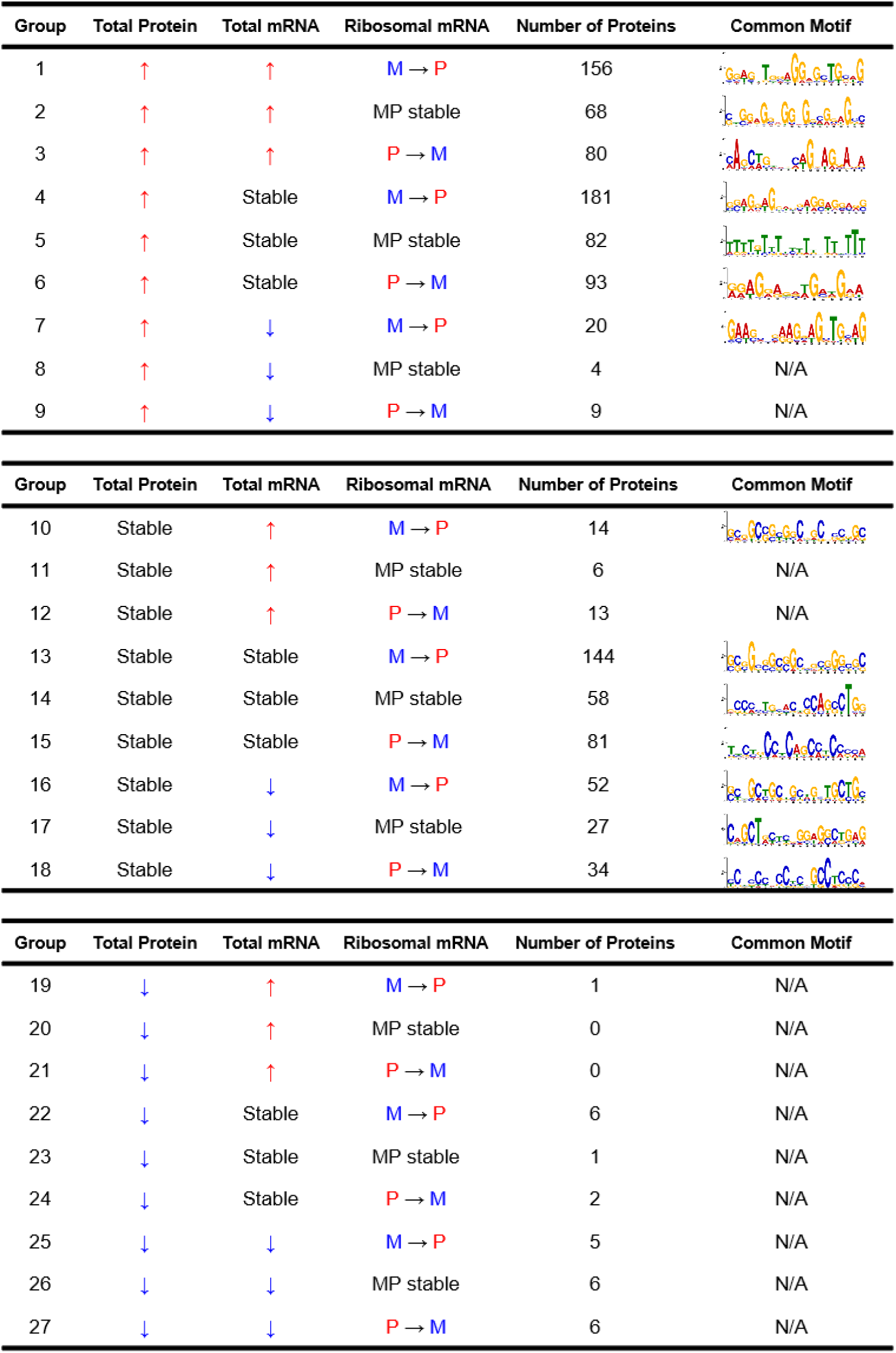
Grouping of proteins and their encoding mRNAs where there was a statistically significant TGF-β1-induced increase (upward arrow), absence of a change, or decrease (downward arrow) for the protein, the total levels of the encoding mRNA, and the polysome/monosome ratio for the mRNA. Common motifs observed in the mRNAs for each group are indicated at right. N/A indicates that there was either no detectable common motif, or too few mRNAs in the group to identify a common motif. M indicates Monosome, P indicates Polysome.

| Group | Total Protein | Total mRNA | Ribosomal mRNA | Number of Proteins | Common Motif |
| --- | --- | --- | --- | --- | --- |
| 1 | ↑ | ↑ | M → P | 156 |  |
| 2 | ↑ | ↑ | MP stable | 68 |  |
| 3 | ↑ | ↑ | P → M | 80 |  |
| 4 | ↑ | Stable | M → P | 181 |  |
| 5 | ↑ | Stable | MP stable | 82 |  |
| 6 | ↑ | Stable | P → M | 93 |  |
| 7 | ↑ | ↓ | M → P | 20 |  |
| 8 | ↑ | ↓ | MP stable | 4 | N/A |
| 9 | ↑ | ↓ | P → M | 9 | N/A |
| 10 | Stable | ↑ | M → P | 14 |  |
| 11 | Stable | ↑ | MP stable | 6 | N/A |
| 12 | Stable | ↑ | P → M | 13 | N/A |
| 13 | Stable | Stable | M → P | 144 |  |
| 14 | Stable | Stable | MP stable | 58 |  |
| 15 | Stable | Stable | P → M | 81 |  |
| 16 | Stable | ↓ | M → P | 52 |  |
| 17 | Stable | ↓ | MP stable | 27 |  |
| 18 | Stable | ↓ | P → M | 34 |  |
| 19 | ↓ | ↑ | M → P | 1 | N/A |
| 20 | ↓ | ↑ | MP stable | 0 | N/A |
| 21 | ↓ | ↑ | P → M | 0 | N/A |
| 22 | ↓ | Stable | M → P | 6 | N/A |
| 23 | ↓ | Stable | MP stable | 1 | N/A |
| 24 | ↓ | Stable | P → M | 2 | N/A |
| 25 | ↓ | ↓ | M → P | 5 | N/A |
| 26 | ↓ | ↓ | MP stable | 6 | N/A |
| 27 | ↓ | ↓ | P → M | 6 | N/A |

### Group 4 mRNAs have a common motif in their coding region

Translation regulation of mRNAs is mediated in general by either microRNAs (Agarwal et al., 2015; Bagnato et al., 2017; Liu et al., 2021), upstream open reading frames (uORFs) (Jewer et al., 2020; Suresh and O’Donnell, 2021; Wethmar et al., 2010) or motifs in the mRNA (Barbieri et al., 2017; Jewer et al., 2020; Ørom et al., 2008; Suresh et al., 2020). Of the 181 proteins in group 4, 152 of the corresponding mRNAs have no known microRNA binding site, and the remaining 29 mRNAs bind a variety of microRNAs, suggesting that the TGF-β1 regulation of translation that we are looking at is not mediated by microRNAs. There were similar percentages of mRNAs with uORFs in 9 representative groups (Figure S2a and S2b), suggesting that the presence of uORFs does not mediate TGF-β1 regulation of translation. A search for common motifs, using an algorithm that looks for both a common sequence and a common structure of the RNA domain (Bailey et al., 2015; Gupta et al., 2007; Moore, 1999), in each of the 22 groups in Table 1 with more than 3 proteins in the group identified common motifs in 14 of the groups. Interestingly, the common motifs of the groups where TGF-β1 affected translation (groups 1, 3, 4, 6, and 7) are mainly located in the coding sequence (CDS) of the mRNAs, while the common motifs in the groups where TGF-β1 had no significant effect on translation (groups 2, 5, 14, and 17) tend to be in the UTRs (Figure S2c). 179 of 181 group 4 mRNAs contain a common motif at p < 10^-6^ (Table 1), and 137 of the 179 had the motif in the CDS. *NEU3* mRNA has two Group 4 motifs in its CDS: 5’-CCTGAAGCCACTGATGGAAG-3’ (315 to 334 in the CDS) and 5’-ACTGAGGCTGGAGGAGGAAG-3’ (1041 to 1060 in the CDS). These results indicate that the common motifs may be correlated to the TGF-β1 induced translation control regulations.

### The motifs in the *NEU3* mRNA are necessary for TGF-β1-increased *NEU3* translation

To test the hypothesis that the group 4 motifs in the *NEU3* mRNA are necessary for TGF-β1 upregulation of NEU3 translation, three Myc-tagged NEU3 expression plasmids were made. Myc-NEU3-WT contains the NEU3 mRNA CDS with a Myc tag inserted at the 3’ end of the protein coding region, with the two group 4 motifs in the *NEU3* CDS. The construct did not contain the *NEU3* 5’ or 3’ UTRs. In the Myc-NEU3-Motif-Mut1 plasmid, the first motif was mutated without causing a frame shift, and NEU3-Motif-Mut2 had an intact motif 1 but a disruption of the second motif, again without causing a frame shift (Supplementary file 7). The plasmids were transfected into human lung fibroblasts, and the cells were treated with or without TGF-β1. We did not observe any differences in the morphology of the cells transfected with any of the plasmids (Figure S3). Staining with anti-Myc antibodies indicated that the Myc tag was not detectable in non-transfected cells (Figures 2a d g and j). TGF-β1 increased levels of Myc-NEU3 protein encoded with a mRNA containing the motifs, but not Myc-NEU3 encoded with a mRNA with a mutation in either motif (Figures 2a d g and j). TGF-β1 did not significantly affect total levels of the *Myc-NEU3* mRNAs (Figure 3a), but did cause wild-type *Myc-NEU3* mRNA to shift from monosomes to polysomes, and this effect was abolished when either of the NEU3 motifs were mutated (Figure 3d). Together this indicates that both of the group 4 motifs in the *NEU3* mRNA are necessary for the TGF-β1-induced *NEU3* translation.

**Figure 2.**
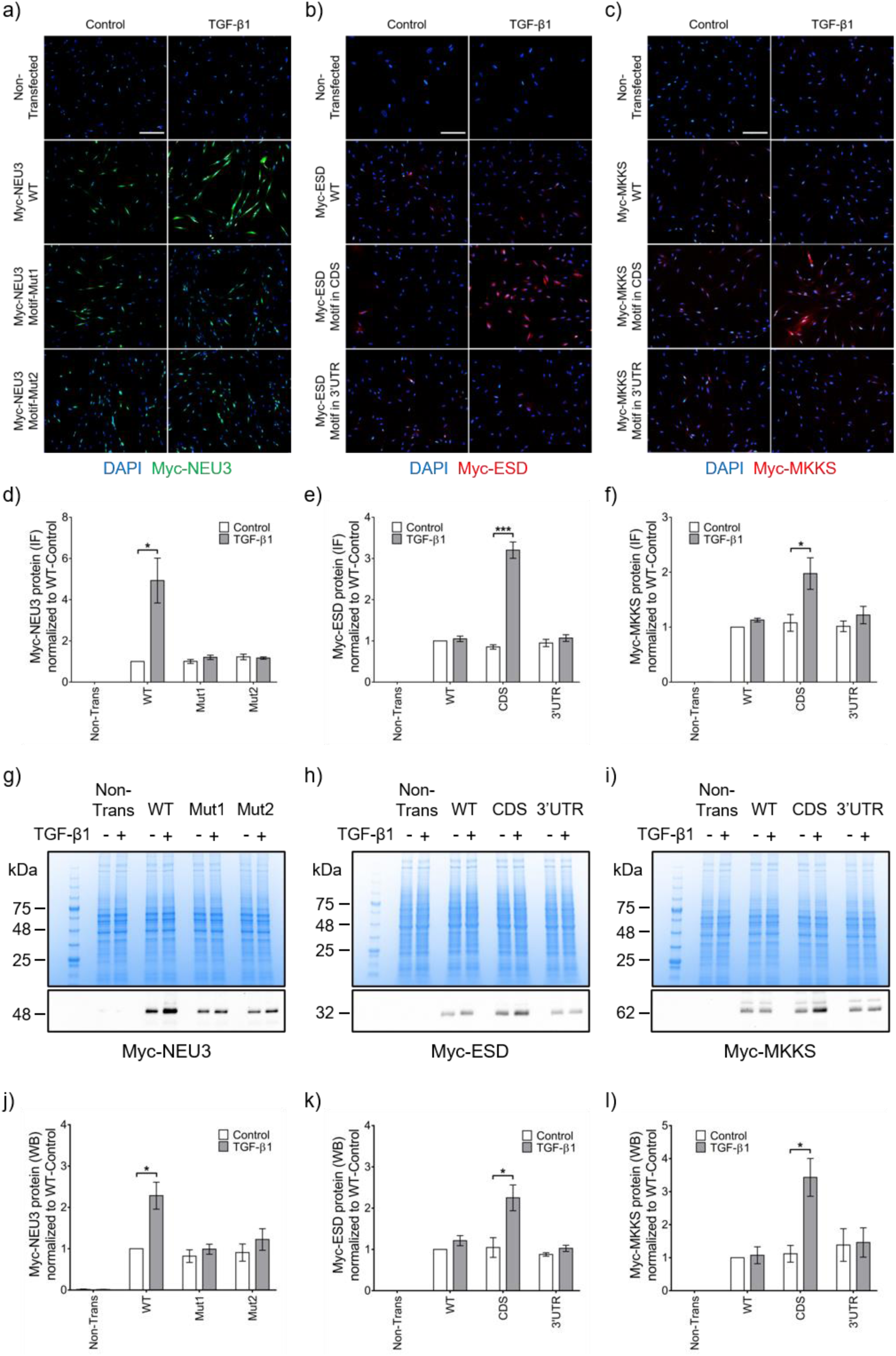
The motifs in the NEU3 mRNA are necessary for TGF-β1-increased NEU3 translation, and the group 4 motif is sufficient for TGF-β1-induced translation of ESD and MKKS. **a)** Human lung fibroblasts transfected with the indicated constructs were cultured without (control) or with TGF-β1 and then stained with anti-Myc-tag antibodies. Green indicates Myc-NEU3 staining and blue is DAPI counterstain. Bar is 100 μm. See Figure S3 for phase images. Images are representative of 3 independent experiments. **b-c)** Human lung fibroblasts transfected with the indicated constructs were cultured without (control) or with TGF-β1 and then stained with anti-Myc-tag antibodies. Red indicates Myc-ESD or Myc-MKKS staining and blue is DAPI counterstain. Images are representative of 3 independent experiments. **d-f)** Quantification of **a-c**. **g)** Myc-NEU3 in transfected and non-transfected HLFs was confirmed by western blotting with an anti-Myc-tag antibody. Top image is Coomassie stained gel of cell lysates, bottom image is western blot. Molecular masses in kDa are at left. **h-i)** Myc-ESD and Myc-MKKS in transfected and non-transfected HLFs were confirmed by western blotting with an anti-Myc-tag antibody. Images in g-i are representative of 3 independent experiments. **j-l)** Quantification of **g-i**. Values in **d-f** and **j-l** are mean ± SEM, n=3. * p < 0.05, *** p < 0.001 (t-tests).

**Figure 3.**
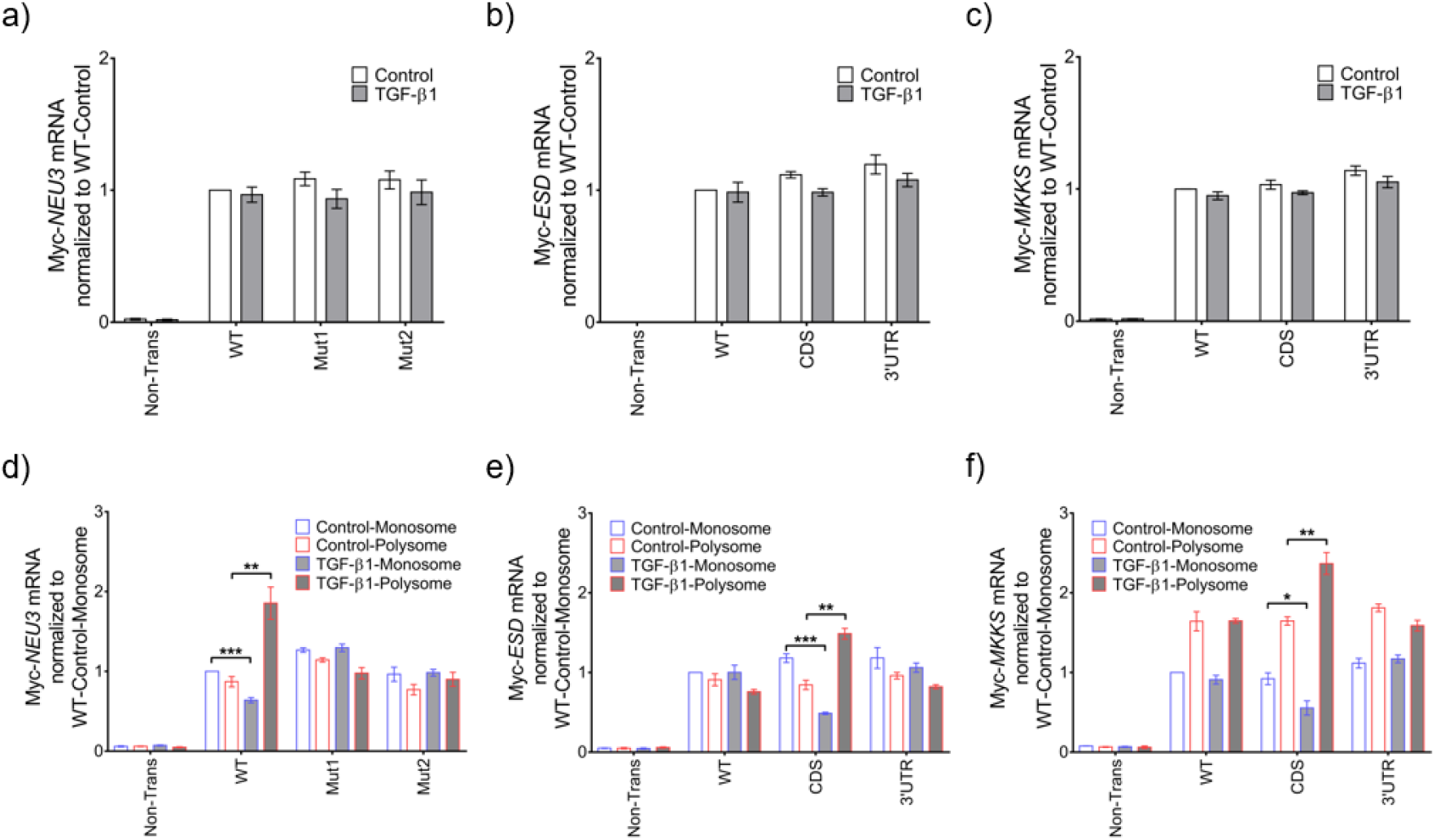
The group 4 motif is necessary and sufficient for TGF-β1-induced monosome/polysome mRNA shifts. **a-c)** Human lung fibroblasts transfected with the indicated constructs were cultured without (control) or with TGF-β1. Total Myc-*NEU3*, Myc-*ESD*, and Myc-*MKKS* mRNA levels were determined by qPCR with primers targeting the junction between the Myc-tag and NEU3, ESD, or MKKS sequences. **d-f)** Ribosomal Myc-*NEU3*, Myc-*ESD*, and Myc-*MKKS* mRNA levels were determined by qPCR with primers targeting the junction between the Myc-tag and NEU3, ESD, or MKKS sequences. Values in a-f are mean ± SEM, n=3. * p < 0.05, ** p < 0.01, *** p < 0.001 (t-tests).

### The group 4 motif is sufficient for TGF-β1-induced translation

To determine if the group 4 motif is sufficient for TGF-β1-induced translation, we examined ESD, a protein from group 14 where TGF-β1 did not significantly affect protein levels, mRNA levels, or mRNA translation. As expected, the group 4 motif was not found in the *ESD* mRNA with MEME. As above, we transformed human lung fibroblasts with constructs encoding Myc-tagged ESD, Myc-tagged ESD encoded by mRNA with the representative group 4 motif (GGAGGAGGAGGAGGAGGAGG) inserted in the CDS without causing a frame shift, and Myc-tagged ESD encoded by mRNA with the group 4 motif inserted in the 3’UTR. We did not observe any differences in the morphology of the cells transfected with any of the plasmids (Figure S3). TGF-β1 had no significant effect on ESD protein levels when the motif was absent, but increased Myc-ESD protein levels when the motif was in the *Myc-ESD* CDS, but not the *Myc-ESD* 3’ UTR (Figures 2b, e, h, and k). TGF-β1 did not affect total levels of the *Myc-ESD* mRNAs (Figure 3b), and did not cause *Myc-ESD* mRNA (with the wild-type *ESD* coding sequence) to shift from monosomes to polysomes (Figure 3e). When the motif was inserted in the ESD coding sequence but not the 3’ UTR, TGF-β1 caused the *Myc-ESD* mRNA to shift from monosomes to polysomes (Figure 3e). Similar results, including an effect of the motif in the CDS but not in the 3’ UTR, were obtained using MKKS, a protein in the same group as ESD (Figures 2c, f, i, and l, and 3c and 3f). Together these results indicate that inserting the group 4 motif in the CDS of a mRNA is sufficient for TGF-β1-induced translation, and that the motif does not potentiate translation when it is in the 3’UTR of a mRNA.

### Changes in polysome/monosome ratios for stable mRNAs can predict changes in proteins in HLF and mouse lungs

To further check the hypothesis that monosome/polysome shifts in mRNAs affect protein levels, we checked 8 protein/ mRNA pairs. Four of these showed monosome to polysome mRNA shifts, and for two of these the protein levels were too low to show up in the proteomics data (Table 2). An additional four showed polysome to monosome mRNA shifts, and two of these had no proteomics reads (Table 2). Protein expression of the 8 mRNAs were checked in TGF-β1 treated HLF cells by staining western blots, and by staining bleomycin treated mouse lung tissues at day 10 (TGF-β1 levels are high in the lungs in the bleomycin mouse model at this time point (Figure 4c and d). TGF-β1 increased levels of three of the four monosome to polysome mRNA shift proteins in HLF (the anti-MTCL1 antibody did not show staining of western blots), and all four showed upregulation in bleomycin-treated mouse lungs (Figure 4a). Similarly, TGF-β1 decreased levels of three of the four polysome to monosome mRNA shift proteins in HLF (the anti-APC4 antibody did not show staining of western blots), and all four showed downregulation in bleomycin-treated mouse lungs (Figure 4e). These results suggest that data from RNA-seq of monosomes and polysomes, in addition to RNA-seq of total RNA, can increase the ability to predict (albeit without 100% certainty) the effects of TGF-β1 on protein levels.

**Figure 4.**
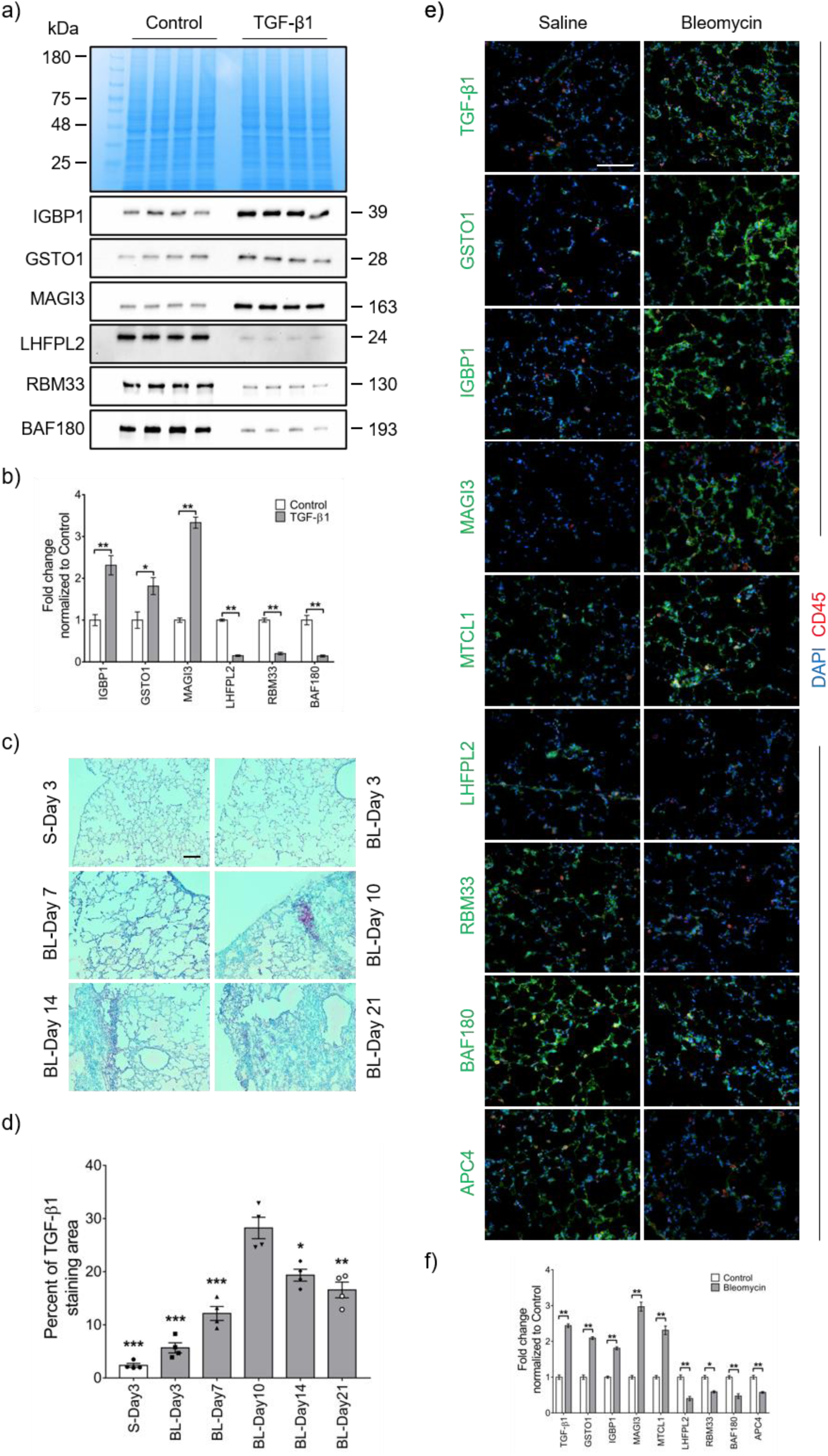
Predictions verified in fibroblasts and mouse lungs. **a)** Human lung fibroblasts were cultured without (control) or with TGF-β1 and then western blots of cell lysates were stained with anti-IGBP1, GSTO1, MAGI3, LHFPL2, RBM33 or BAF180 antibodies. Top image is Coomassie stained gel of cell lysates, bottom image is western blot. Molecular masses in kDa are at left for the Coomassie gel and at right for the western blots. Pairs of columns (control and TGF-β1) are independent experiments. **b)** Quantification of **a**. Values are mean ± SEM, n = 4. * p < 0.05, ** p < 0.01 (t-tests). **c)** Mouse lung tissues collected at day 3, 7, 10, 14 and 21 after saline (S) or bleomycin (BL) treatment were stained with anti-TGF-β1 antibodies. Red indicates TGF-β1 positive stain. Bar is 100 μm. Images are representative of 4 independent experiments (3 male mice and 1 female mouse). **d)** Quantification of **c**. Values are mean ± SEM, n = 4. * p < 0.05, ** p < 0.01, *** p < 0.001 vs BL-Day 10 (1-way ANOVA, Dunnett’s test). **e)** Mouse lung tissues collected at day 10 after saline or bleomycin treatment were stained with anti-TGF-β1, GSTO1, IGBP1, MAGI3, MTCL1, LHFPL2, RBM33, BAF180, or APC4 antibodies and anti-CD45 antibodies. Red is CD45 positive stain, green is positive stain for the indicated protein, and blue is DAPI staining of nuclei. Bar is 100 μm. Images are representative of 4 independent experiments (3 male and 1 female mice). **f)** quantification of **e**. Values are mean ± SEM, n = 4. * p < 0.05, ** p < 0.01 (t-tests).

**Table 2.** Summary of the results from Figure 4. An upward arrow indicates a TGF-β1-induced or bleomycin-induced increase, a downward arrow indicates a decrease. Ab N/A for WB indicates the antibody was unable to detect the protein on western blots of lysates from human lung fibroblasts. M indicates Monosome, P indicates Polysome.

| ID | Protein | Total mRNA | Mono / Poly mRNA | Bleo mice | HLF cell line |
| --- | --- | --- | --- | --- | --- |
| IGBP1 | ↑ | ↓ | M → P shift | ↑ | ↑ |
| GSTO1 | ↑ | ↓ | M → P shift | ↑ | ↑ |
| MAGI3 | No proteomics reads | Stable | M → P shift | ↑ | ↑ |
| MTCL1 | No proteomics reads | Stable | M → P shift | ↑ | Ab N/A for WB |
| LHFPL2 | ↓ | Stable | P → M shift | ↓ | ↓ |
| RBM33 | ↓ | Stable | P → M shift | ↓ | ↓ |
| BAF180 | No proteomics reads | Stable | P → M shift | ↓ | ↓ |
| APC4 | No proteomics reads | Stable | P → M shift | ↓ | Ab N/A for WB |

### TGF-β1 regulates group 4 motif interacting RNA-binding proteins, and one of the RBPs, DDX3, increases in human IPF lung

UV cross-linking and immunoprecipitation (CLIP) identifies the RNAs in a cell line or tissue that bind to a specific RNA-binding protein (Krakau et al., 2017; Moore et al., 2014; Shah et al., 2017; Weyn-Vanhentenryck et al., 2014). To elucidate how TGF-β1 induces the motif-dependent translation of group 4 mRNAs, published CLIP-seq databases (Hu et al., 2017; Yang et al., 2015; Zhao et al., 2022; Zhu et al., 2019) were analyzed to identify RNA-binding proteins that bind group 4 mRNAs. Many RNA-binding proteins that bind to group 4 mRNAs were found, and 17 of them bind to more than 50% of the 182 group 4 mRNAs (Figure 5a and b). Since some group 4 mRNAs may not be expressed in the cells and tissues used for the CLIP-seq databases, a higher percentage of the group 4 mRNAs may bind the proteins in Figure 5a and b, and the values in Figure 5a and b are thus a low estimate. To select the group 4-motif specific RNA-binding proteins, an interaction prediction between the group 4-motif and the 17 RNA-binding proteins was done using RPIseq (Muppirala et al., 2011; Muppirala et al., 2013) (Figure 5c). For further analysis, we chose AGO2, UPF1, DDX3, U2AF2 and CELF2, as these are known translation regulators (Hurt et al., 2013; Karginov and Hannon, 2013; Lee et al., 2008; Mallory et al., 2020; Palangat et al., 2019) and showed high interaction probabilities with the Group 4 motif. AGO2 and UPF1 are P-body proteins (Hubstenberger et al., 2017), and phosphorylation of AGO2 and UPF1 leads to mRNA translational repression (Durand et al., 2016; Horman et al., 2013; Isken et al., 2008). DDX3 is a translation stimulator (Chen et al., 2018; Soto-Rifo et al., 2012; Soto-Rifo et al., 2013). U2AF2 plays an important role in RNA splicing (Laliotis et al., 2021; Maji et al., 2020). CELF2 is a translation suppressor (Mallory et al., 2020; New et al., 2019; Yeung et al., 2020). ALYREF, a RBP that binds relatively few group 4 mRNAs (Figure 5a and 5b), and which has a low predicted binding to the group 4 motif (Figure 5c) was chosen as a negative control.

**Figure 5.**
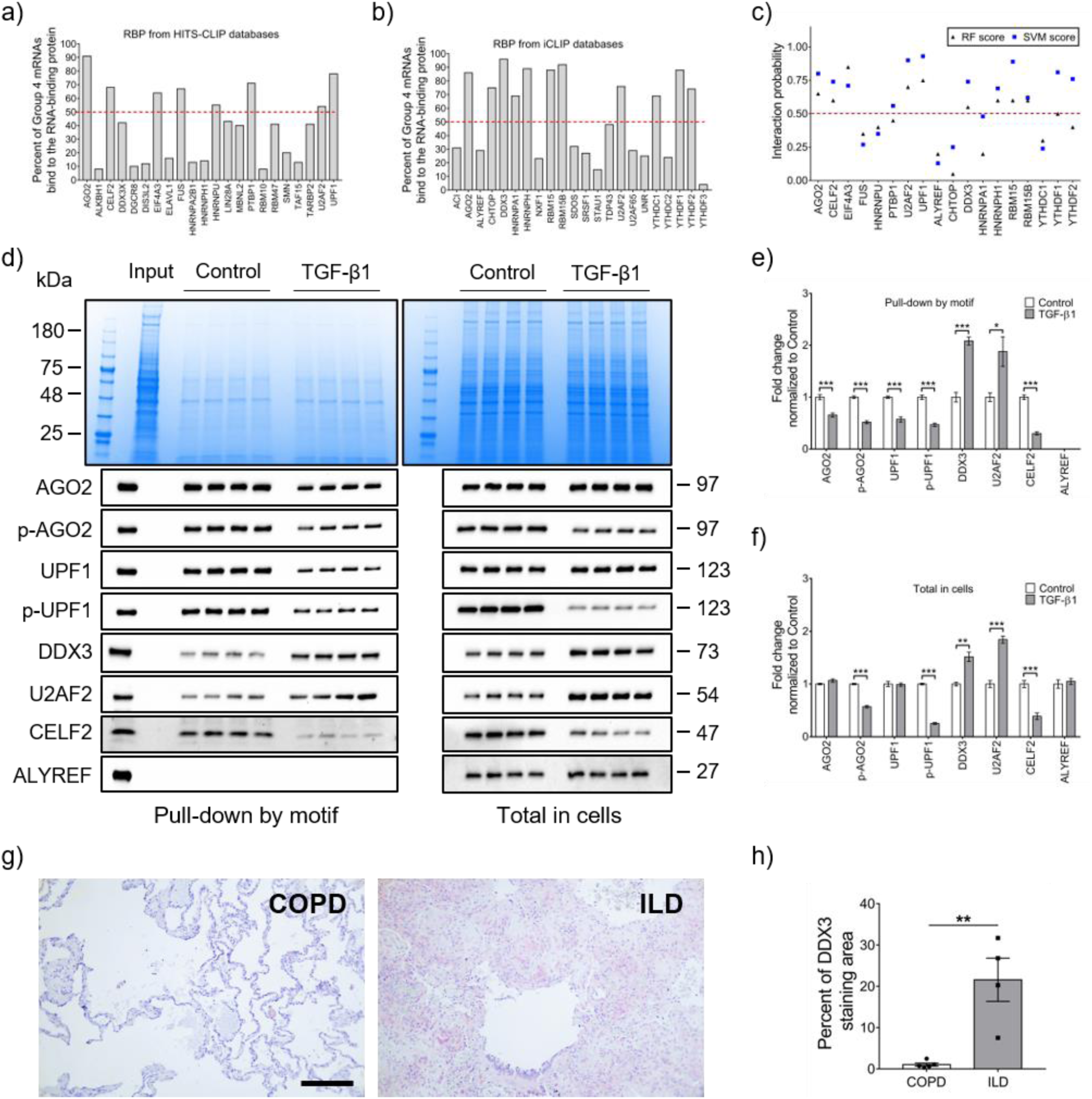
TGF-β1 regulates group 4 motif interacting RNA-binding proteins, and one of the RBPs, DDX3, is upregulated in human IPF lung. **a-b)** RNA-binding proteins that can bind to group 4 mRNAs were identified from the HITS-CLIP database and the iCLIP database. Values are the percent of the Group 4 mRNAs that were predicted to bind to the indicated RNA-binding protein using HITS-CLIP (a) or iCLIP (b). Red dashed line indicates 50%. **c)** Group 4 motif specific RNA-binding proteins were selected by RNA-protein interaction prediction software (RPISeq). The interaction probability was predicted and scored by a Random Forest classifier (triangles) and a Support Vector Machine classifier (squares). RNA-binding proteins with scores higher than 0.5 (red dashed line) were considered as having a high possibility of interacting with the indicated RNA-binding protein. **d) Left:** The biotin-tagged RNA motif was used to pull down material from control and TGF-β1-treated cells. The Coomassie-stained gel shows the material in the pull-down samples, and the Western blots show staining of the samples with the indicated antibodies. **Right:** The Coomassie-stained gel shows total cell lysate, and the Western blots show staining of the total cell lysate with the indicated antibodies. For Left and Right, pairs of columns (control and TGF-β1) are independent experiments. Molecular masses in kDa are at left for the Coomassie stained gel and at right for the western blots. **e-f)** Quantification of protein levels in pull-down samples and total samples. Values in **e** and **f** are mean ± SEM, n=4. * p < 0.05, ** p < 0.01, *** p < 0.001 (t-tests). **g)** Lung tissue sections from COPD or IPF patients were stained with anti-DDX3 antibodies. **Left:** Section from a COPD patient with FVC >80%. **Right:** Section from an IPF patient with FVC <50%. Positive staining was identified by red color, and nuclei are counterstained blue. Bar is 200 μm. Images are representative of 4 patients per group. **h)** The percentage area of lung tissue stained by DDX3 antibody. Values are mean ± SEM, n = 4. ** p < 0.01 (t-test).

To determine if the interactions of these proteins with the motif is regulated by TGF-β1, 5’ biotin-labeled GGAGGAGGAGGAGGAGGAGG RNA (the group 4 consensus motif) was used for pull-down assays with streptavidin-conjugated beads from lysates of the four different human lung fibroblast cell lines cultured with or without TGF-β1. Using antibodies against all of the above RNA-binding proteins, and antibodies against phosphorylated AGO2 and UPF1, we observed that TGF-β1 decreased the interaction of the tagged motif with AGO2, phospho-AGO2, UPF1, phospho-UPF1, and CELF2, and increased the motif binding to DDX2 and U2AF2 (Figure 5d and e). The software predicted that the RNA-binding protein ALYREF would not bind the motif, and we indeed observed that the tagged motif did not pull down this protein (Figure 5d and e). Using the antibodies to stain western blots of whole cell lysates, we observed that TGF-β1 did not significantly change levels of AGO2, UPF1, or ALYREF, decreased levels of phospho-AGO2, phospho-UPF1, and CELF2, and increased levels of DDX3 and U2AF2 (Figure 5d and f). The *DDX3* mRNA has a group 4 motif, and is in group 4, suggesting that DDX3 may potentiate its own upregulation in response to TGF-β1. These results suggest that TGF-β1 regulates at least 5 RNA-binding proteins, some of which may in turn regulate *NEU3* translation.

To determine if DDX3 is associated with human lung fibrosis, we examined the distribution of DDX3 in lung tissue from chronic obstructive pulmonary disease (COPD) patients with relatively normal lungs (> 80% Forced vital capacity; FVC) and idiopathic pulmonary fibrosis (IPF) patients with advanced disease (<50% FVC). Lung tissue from COPD patients had limited DDX3 staining except for occasional single positive cells in the alveolar space (Figure 5g). In the lung tissue from pulmonary fibrosis patients, DDX3 was distributed throughout the tissue, including the lung epithelium, leukocytes, and fibrotic areas (Figure 5g). The IPF lungs had a greater area of tissue DDX3 staining than the COPD lungs (Figure 5h). Together, these data indicate that the levels of DDX3 are increased in human lung fibrosis.

### DDX3 siRNA and DDX3 inhibitors block TGF-β1 induced NEU3 expression in HLF

To determine if DDX3 mediates the effect of TGF-β1 on NEU3 translation, HLFs were transfected with DDX3 siRNA or negative control siRNA. 24 hours later, HLFs were treated with or without 10 ng/ml TGF-β1. After 48 hours, the culture medium was removed and cells were lysed, collected, and assessed by western blotting. Compared to non-transfected cells, DDX3 siRNA decreased DDX3 and NEU3 (Figure 6a-c). As shown above, TGF-β1 increased DDX3 and NEU3 levels, and DDX3 siRNA blocked the TGF-β1 induced DDX3 and NEU3 upregulation. Negative control siRNA did not decrease DDX3 or NEU3 (Figure 6a-c). We also determined if two DDX3 inhibitors (RK-33 and IN-1) (Brai et al., 2016; Tantravedi et al., 2019) could reduce TGF-β1 mediated NEU3 levels. RK-33 at 10 µM inhibits DDX3 in medulloblastoma and breast cancer cell lines (Heerma van Voss et al., 2018; Tantravedi et al., 2019). IN-1 at 50 µM inhibits DDX3 in PBMCs and hepatocellular carcinoma cells (Brai et al., 2016). Compared to control, TGF-β1 increased NEU3 levels (Figure 6d). Both RK-33 and IN-1 inhibited the up-regulation of NEU3 induced by TGF-β1 (Figure 6d). Both inhibitors also reduced NEU3 in cells cultured in the absence of TGF-β1 (Figure 6d). These data indicate that DDX3 mediates the effect of TGF-β1 on NEU3.

**Figure 6.**
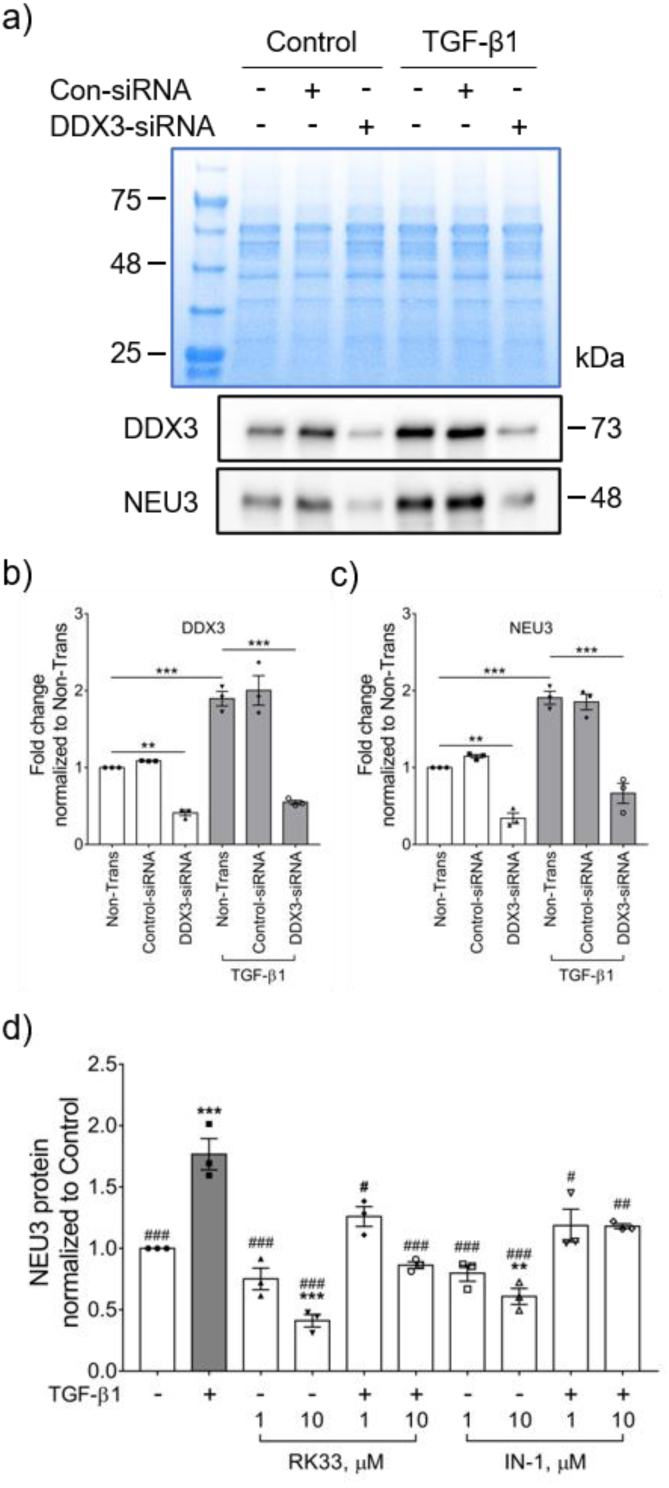
DDX3 siRNA and DDX3 inhibitors block TGF-β1-increased NEU3 levels. **a)** Human lung fibroblasts were transfected with control or DDX3 siRNA, and cultured without (control) or with TGF-β1 and then stained with anti-DDX3 and NEU3 antibodies. Protein levels were determined by western blotting. Top image is a Coomassie stained gel of cell lysates, bottom images are western blots. Images are representative of 3 independent experiments. Molecular masses in kDa are at left for the Coomassie stained gel and at right for western blots. **b-c)** Quantification of **a**. **d)** Human lung fibroblasts were treated with DDX3 inhibitors, and cultured without (control) or with TGF-β1 and then stained with anti-NEU3 antibody. Protein levels were determined by western blotting. In b-d, values are mean ± SEM, n = 3. ** p < 0.01, *** p < 0.001 vs control, # p < 0.05, ## p < 0.01, ### p < 0.001 vs TGF-β1 treated (1-way ANOVA, Dunnett’s test).

### RK-33 attenuates pulmonary fibrosis in mice

Genetic deletion of NEU3 or inhibiting NEU3 inhibits pulmonary fibrosis in mice (Karhadkar et al., 2020a; Karhadkar et al., 2021; Karhadkar et al., 2017), and as described above, the DDX3 inhibitor inhibits TGF-β1-induced upregulation of NEU3 in lung fibroblasts. To determine if RK-33 might inhibit bleomycin-induced pulmonary fibrosis in mice, C57BL/6 mice were treated with an oropharyngeal aspiration of saline or bleomycin, and then starting 10 days after saline or bleomycin, were given intraperitoneal injections of 20 mg/kg RK-33 or buffer control every 2 days. Compared to bleomycin-treated mice, mice that received bleomycin and were then treated with RK-33 had a higher survival rate (Figure 7a). In saline-treated mice, RK-33 did not affect survival. As previously observed, compared with saline treated mice, bleomycin-treated mice lost weight over the first 10 days, and did not recover their initial weight by 21 days (Figure 7b). In saline treated mice, RK-33 did not significantly affect body weights. RK-33 attenuated the bleomycin-induced weight loss from days 17-21 (Figure 7b).

**Figure 7.**
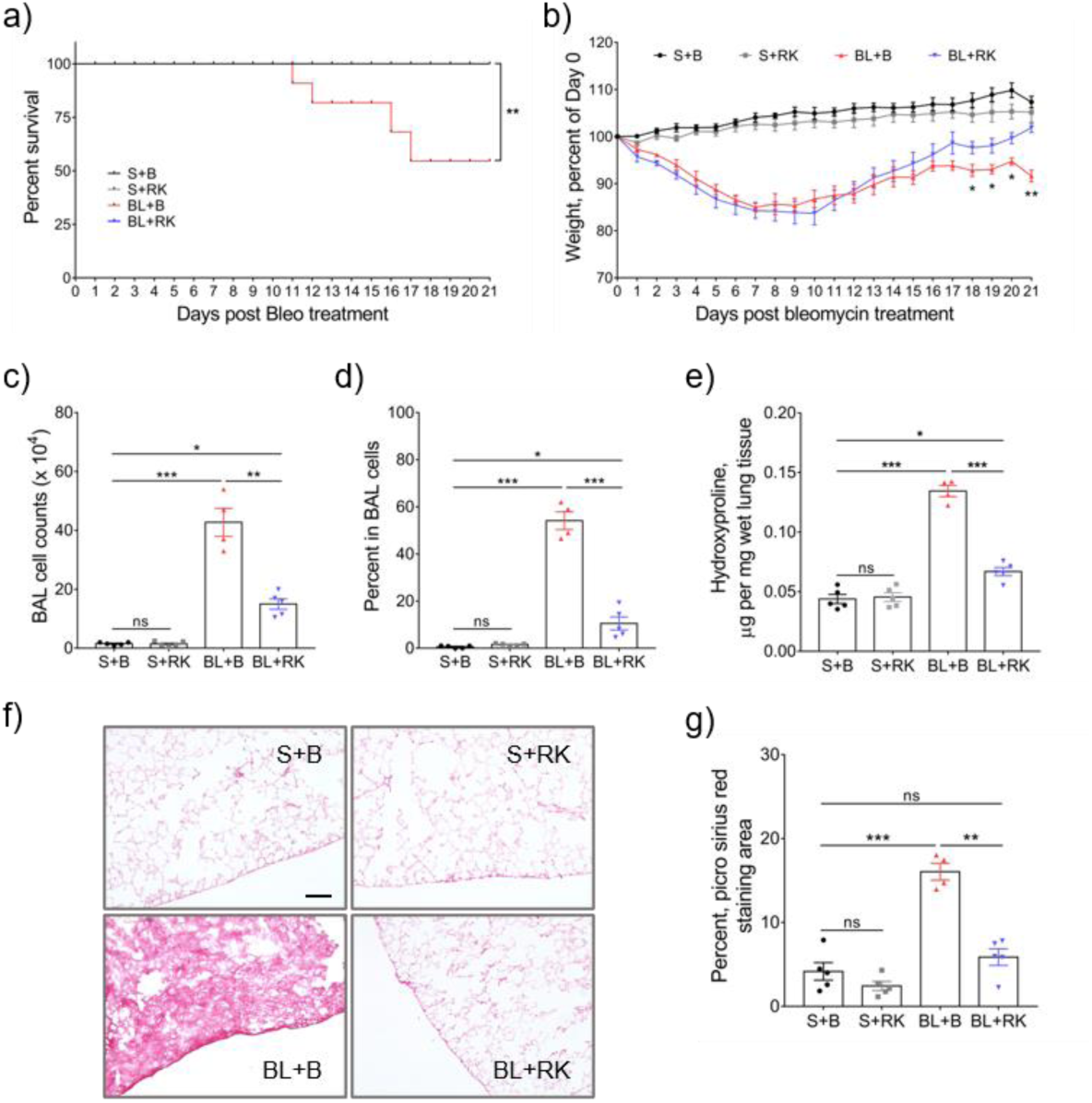
RK-33 attenuates pulmonary fibrosis in mice. Mice were treated with saline or bleomycin, and then starting at day 10 after saline or bleomycin, mice were given 20 mg/kg RK-33 or buffer control every 2 days. Saline with buffer (S+B), saline with RK-33 (S+RK), bleomycin with buffer (BL+B), and bleomycin with RK-33 (BL+RK). **a)** Survival of mice. n = 5 mice for S+B, S+RK, and BL+RK. n = 8 mice for BL+B. ** p < 0.01 vs BL+B (Mantel–Cox test). **b)** Percent change in body weight. Values are means ± SEM, n = 5 mice for S+B, S+RK, and BL+RK, n = 4 mice that survived to day 21 for BL+B. * p < 0.05, ** p < 0.01, BL+B vs BL+RK (t-test). **c)** Total number of cells detected in mouse bronchoalveolar lavage fluid (BALF). **d)** Percent of lymphocytes in the BAL cell population as assessed by Wright-Giemsa staining. **e)** Hydroxyproline levels in mouse lung. **f)** PicroSirius-red stained mouse lung sections. Bar is 100 μm. Images are representative of 5 mice for S+B, S+RK, and BL+RK, and 4 surviving mice for BL+B. **g)** Quantification of PicroSirius-red positive stain area in 3 randomly chosen areas of each mouse lung. For **c-e** and **g**, values are mean ± SEM, n = 5 mice for S+B, S+RK, and BL+RK, n = 4 mice for BL+B. ns = non-significant, * p < 0.05, ** p < 0.01, *** p < 0.001 (1-way ANOVA, Dunnett’s test).

Bleomycin upregulates inflammatory cell counts in mouse lung (Izbicki et al., 2002; Karhadkar et al., 2017; Swaney et al., 2010). Saline-treated control mice and saline-treated mice that received RK-33 showed similar BAL cell counts and percent of lymphocytes in the BAL (Figure 7c and d). Compared to saline treated mice, bleomycin treated control mice showed increased BAL cell counts with an increased percentage of lymphocytes (Figure 7c and d). However, compared to the bleomycin-control mice, bleomycin-treated and RK-33 injected mice had significantly less BAL and lymphocytes.

To determine whether RK-33 injections can decrease bleomycin-induced fibrosis, lung sections were stained with picrosirius red to detect collagen, and hydroxyproline levels were measured in the lung tissue lysates. Saline-treated control mice and saline-treated mice that received RK-33 showed similar hydroxyproline levels and picrosirius red staining (Figures 7e-7g). Compared to saline treated mice, bleomycin-treated mice had increased levels of hydroxyproline and increased lung tissue staining for picrosirius red (Figures 7e-7g). Compared to the bleomycin-control mice, bleomycin-treated and RK-33 injected mice had reduced levels of hydroxyproline and reduced lung tissue staining for picrosirius. Together, these results indicate that RK-33 can attenuate lung fibrosis in mice.

Bleomycin aspiration also causes an increase in the number of inflammatory cells and an increase in the levels of TGF-β1 and NEU3 in lung tissue (Karhadkar et al., 2020a; Karhadkar et al., 2021; Karhadkar et al., 2017). Compared to saline treated mice, bleomycin increased the number of CD45+, CD11b+, and CD11c+ cells remaining in the lungs after BAL (Figures 8a-d). RK-33 injections did not affect the counts of CD45+, CD11b+, or CD11c+ cells in mice treated with saline at day 0, but RK-33 significantly reduced the counts of these cells after bleomycin (Figure 8a-d).

**Figure 8.**
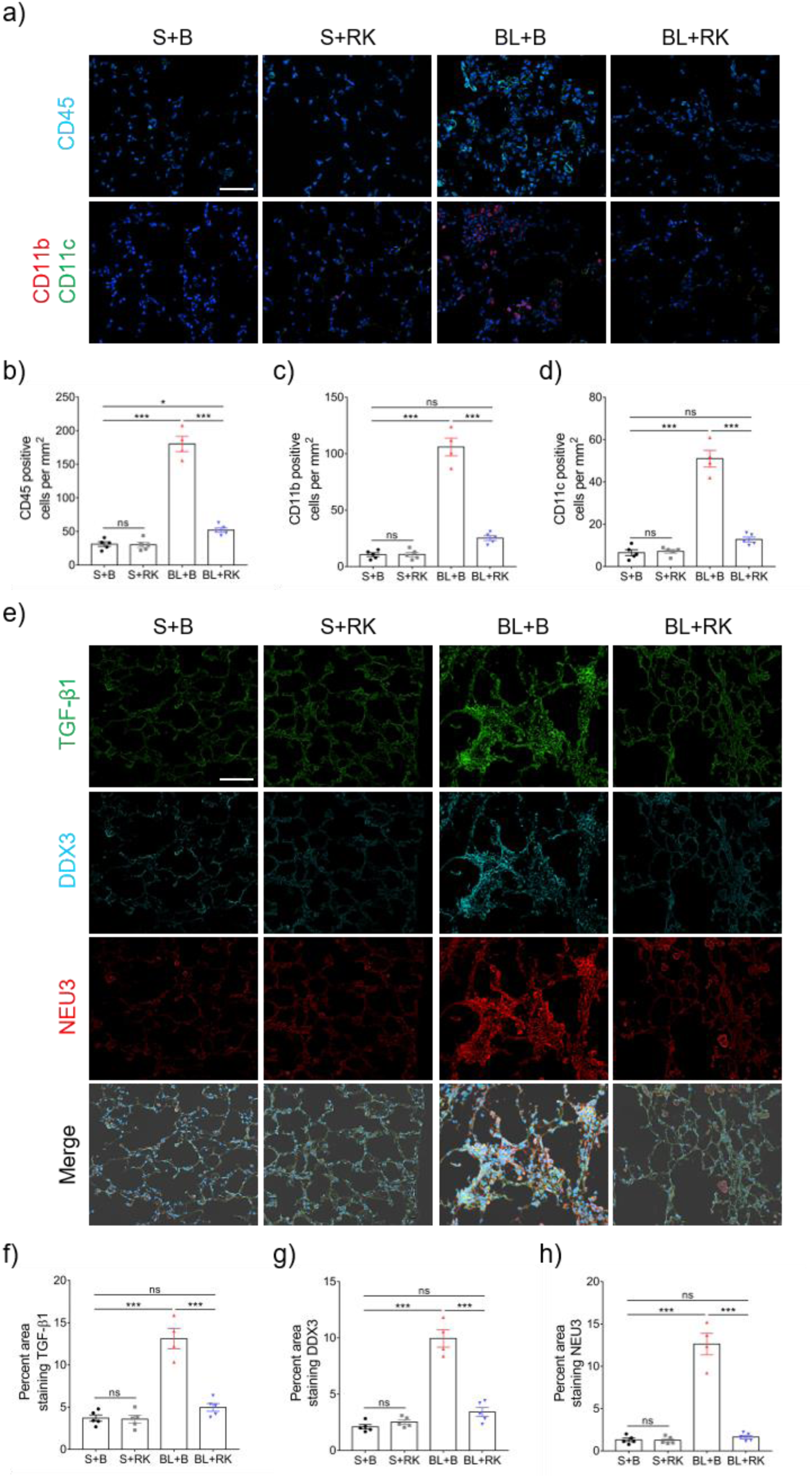
RK-33 reduces bleomycin increased CD45, CD11b, CD11c, TGF-β1, DDX3 and NEU3. **a)** Cryosections of mouse lungs from the Figure 7 experiment were stained with anti-CD45, CD11b, or CD11c antibodies. Cyan indicates CD45, red indicates CD11b, and green indicates CD11c positive stain. Nuclei are stained blue with DAPI. Bar is 100 μm. Images are representative of n = 5 mice for S+B, S+RK, and BL+RK, n = 4 mice for BL+B. **b-d)** Quantification of positive stained cells. **e)** Cryosections of mouse lungs from the Figure 7 experiment were stained with anti-TGF-β1, DDX3, and NEU3 antibodies. Cyan indicates DDX3, red indicates NEU3, and green indicates TGF-β1 positive stain. Nuclei are stained blue with DAPI. Bar is 100 μm. Images are representative of n = 5 mice for S+B, S+RK, and BL+RK, n = 4 mice for BL+B. **f-h)** Quantification of positive stained area in 3 randomly stained field of view for each section from each mouse. Values are mean ± SEM. ns = not significant, *** p < 0.001 (1-way ANOVA, Dunnett’s test).

Compared to saline treated mice, bleomycin increased the expression of TGF-β1, DDX3, and NEU3 in the lung tissue after BAL (Figures 8e-8h). RK-33 injections significantly reduced the levels of all three proteins in lung tissue after bleomycin treatment (Figure 8e-8h). The saline-treated control mice and saline-treated mice that received RK-33 showed similar levels of TGF- β1, DDX3, and NEU3 staining (Figures 8e-8h). Together, these data indicate that RK-33 may reduce bleomycin-induced inflammation, fibrosis, and TGF-β1, DDX3, and NEU3 upregulation in mouse lungs.

## Discussion

In both bacteria and eukaryotes, typically only ∼40% of changes in levels of proteins can be explained by a corresponding change in levels of the associated mRNA (de Sousa Abreu et al., 2009; Maier et al., 2009; Schwanhäusser et al., 2011), indicating that regulating mRNA translation and/or protein degradation plays a more significant role in regulating protein levels than does regulating mRNA levels. In agreement with these observations, we observed that a large fraction of TGF-β1-induced changes in levels within a set of 1,149 proteins in human lung fibroblasts cannot be explained by a corresponding change in the levels of the mRNA encoding the protein. In addition to NEU3, there are at least 181 other proteins in the group 4 subset where TGF-β1 shows a statistically significant absence of a change in mRNA levels, but increases translation of the mRNA and increases levels of the protein. Although in lung epithelial cells TGF-β1 decreases NEU3 degradation in addition to increasing *NEU3* translation (Chen et al., 2020), at least some of the TGF-β1-induced increases in group 4 proteins may be due to increased translation of their mRNAs. In addition, the data indicate that TGF-β1 increases levels of NEU3 in at least two different lung cell types by increasing *NEU3* translation without changing levels of *NEU3* mRNA.

In addition to *NEU3*, all but 2 of the Group 4 mRNAs contain a common 20-nucleotide motif. Altering the motif in *NEU3*, and adding the motif to two different group 14 mRNAs (where TGF-β1 shows a statistically significant absence of an effect on mRNA, translation, and protein levels) indicated that the motif is necessary and sufficient for TGF-β1 regulation of *NEU3* translation. The absence of the motif from two of the group 4 mRNAs suggests that either the software analysis was too stringent and did not detect a group 4-like motif in these 2 mRNAs, or that there is some additional mechanism that mediates TGF-β1-induced increased translation of these mRNAs.

Gene expression in eukaryotes is extensively controlled at the post-transcriptional level by RNA-binding proteins (RBPs) and ribonucleoprotein complexes modulating the stability, transport, editing, and translation of mRNAs (Hafner et al., 2010; Schwanhäusser et al., 2011). We observed that at least 5 RBPs bind the group 4 consensus motif, and all 5 show a regulation by TGF-β1. Argonaute 2 (AGO2) and Regulator of nonsense transcripts 1 (UPF1) are known P-body proteins (Hubstenberger et al., 2017), and the phosphorylation of AGO2 and UPF1 leads to mRNA translational repression (Durand et al., 2016; Horman et al., 2013; Isken et al., 2008). TGF-β1 had no significant effect on levels of these two RBPs in cells, but rather induced their dephosphorylation and caused them to decrease their interaction with the group 4 motif. CUGBP Elav-like family member 2 (CELF2) is a translation suppressor (Mallory et al., 2020; New et al., 2019; Yeung et al., 2020), and TGF-β1 decreased levels of this protein in cells, and possibly as a result decreased the amount of binding of this protein to the group 4 motif. Together, these results suggest a scenario where three different RNA-binding proteins bind to, and inhibit translation of, group 4 mRNAs. In this scenario, TGF-β1 causes these three translation-inhibiting RBPs to decrease their binding to group 4 mRNAs (for two of the proteins, apparently by inducing a dephosphorylation of the RBPs), allowing the group 4 mRNAs to increase their interaction with, or ability to be translated by, ribosomes.

The ATP-dependent RNA helicase DDX3 is a translation stimulator (Chen et al., 2018; Soto-Rifo et al., 2012; Soto-Rifo et al., 2013), and the splicing factor U2AF 65 kDa subunit (U2AF2) plays a role in RNA splicing (Laliotis et al., 2021; Maji et al., 2020). TGF-β1 increased levels of both of these two RBPs in cells, and possibly as a result increased the amount of these two RBPs binding to a tagged group 4 consensus motif. In the above scenario, the TGF-β1-induced increase in levels of the two RBPs, one of which is a known translation stimulator, would increase their ability to compete with the three translation-inhibiting RBPs for binding to the group 4 motif, and the net result would be a TGF-β1-induced increase in translation of group 4 mRNAs. As with the regulation of cyclin-dependent kinases (Arif et al., 2011; Doonan and Kitsios, 2009; Kalous et al., 2020), having multiple positive and negative regulators would allow multiple signals and events in a cell to regulate group 4 mRNA translation with specific dominance relationships.

*NEU3* mRNA contains two of the group 4 motifs, and altering either of these abolished the ability of TGF-β1 to increase *NEU3* translation. This suggests that for NEU3, there is an unknown interaction of the motifs, and/or the proteins that bind the motifs, such that the presence of both motifs is necessary for the ability of TGF-β1 to increase *NEU3* translation.

Due to the limitations of the proteomics, we were only able to examine the regulation of the more prevalent proteins in human lung fibroblasts. As others have found in other systems, RNA-seq of total RNA (de Sousa Abreu et al., 2009; Maier et al., 2009; Schwanhäusser et al., 2011) is a poor predictor of whether a specific signal, in this case TGF-β1, causes an increase or decrease of a protein encoded by a mRNA. Fractionation of mRNA into monosomes and polysomes is also a poor predictor, as exemplified by the 144 proteins in group 13 where there is a clear increase in translation, but no increase in protein levels. For unknown reasons, the combination of a group 4 motif and a monosome to polysome shift is a good predictor of an increase in protein levels, at least within the test group, and an intriguing possibility is that combining motif analysis with RNA-seq of monosomes and polysomes may increase the ability to predict the effects of a signal such as TGF-β1 on protein levels in a situation where there is no antibody available to detect the protein, and the protein levels are too low to be detected by proteomics.

In the bleomycin model of pulmonary fibrosis, treatment with the DDX3 inhibitor RK-33 potentiated survival, attenuated inflammation and fibrosis, and reduced lung tissue levels of DDX3, TGF-β1, and NEU3 in young male mice. Further work is thus needed to determine if this effect occurs in young female mice and old mice. DDX3 was also increased in human fibrotic lung tissue, suggesting that high levels of DDX3 may be associated with disease progression in IPF. Increased levels of DDX3 correlate with poor prognosis in many tumors, and treatment with RK-33 reduces tumor growth and improves survival (Bol et al., 2015; Kukhanova et al., 2020; Tantravedi et al., 2019). Possibly as a result of regulating RNA translation, DDX3 also inhibits apoptosis and promotes cell proliferation, adhesion, and motility, all factors that also promote lung inflammation and fibrosis (Mo et al., 2021; Wolters et al., 2014). These reports and our studies suggest that targeting DDX3, and possibly other RNA-binding proteins, with inhibitors such as RK-33 to inhibit upregulation of the translation of specific mRNAs may be useful as a possible therapeutic for lung inflammation and fibrosis.

## Acknowledgments

We thank the LARR (Laboratory Animal Resources and Research) staff at Texas A&M University for animal care.

We thank Travis Mosley and Shiyu Jing for preliminary analysis of miRNA and uORF datasets, and Issam Ismail for testing some of the antibodies for western blotting.

We thank Dr. Isabella Farhy and Dr. Jennifer Dulin, Texas A&M University, for use of their cryostats.

We are grateful to Dr. Carol Feghali-Bostwick, Medical University of South Carolina, for the human lung fibroblast (HLF) cell lines,

This work was supported by National Institutes of Health National Heart, Lung, and Blood Institute [Grant R01 HL132919].

## Author contributions

Conceptualization, W.C. and R.H.G.; Methodology, W.C. and R.H.G.; Investigation, W.C. and D.P.; Writing-Original Draft, W.C. and R.H.G.; Writing-Review & Editing, W.C., D.P. and R.H.G.; Funding Acquisition, R.H.G.

---

**Figure S1.**
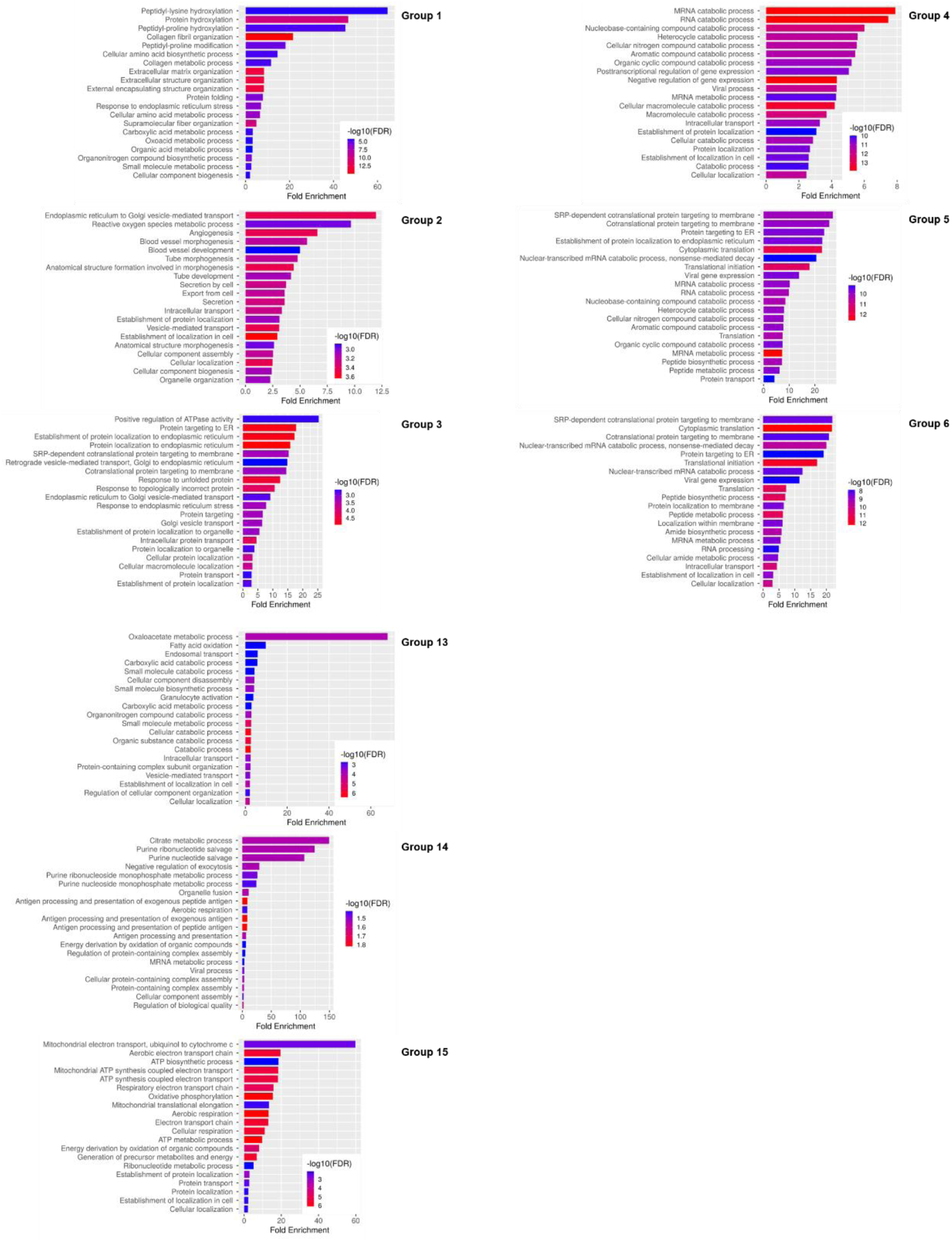
Gene ontology (GO) biological process analysis of groups with 20 or more proteins. Significance (p < 0.05) was determined by Fisher’s exact test with False Discovery Rate (FDR) correction.

**Figure S2.**
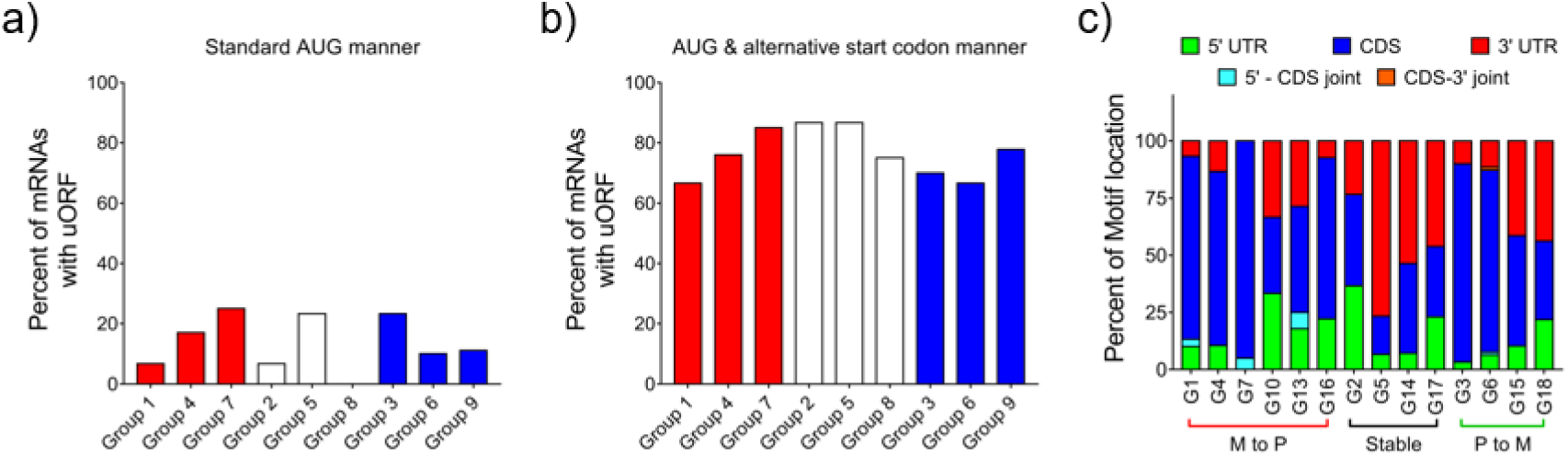
Percent of mRNAs with an upstream open reading frame (uORF) and locations of motifs in mRNAs. **a-b)** Each mRNA in the indicated group was assessed for an uORF with ether an AUG near the 5’ end of the ORF or an AUG or alternative start codon near the 5’ end of the uORF. **c)** Motif location was analyzed in mRNAs which showed a TGF-β1-induced monosome to polysome (M to P) shift (groups 1, 4, 7, 10, 13, 16), mRNAs which showed a polysome to monosome (P to M) shift (groups 3, 6, 15, 18), and mRNAs which showed no change with TGF-β1 treatment (stable).

**Figure S3.**
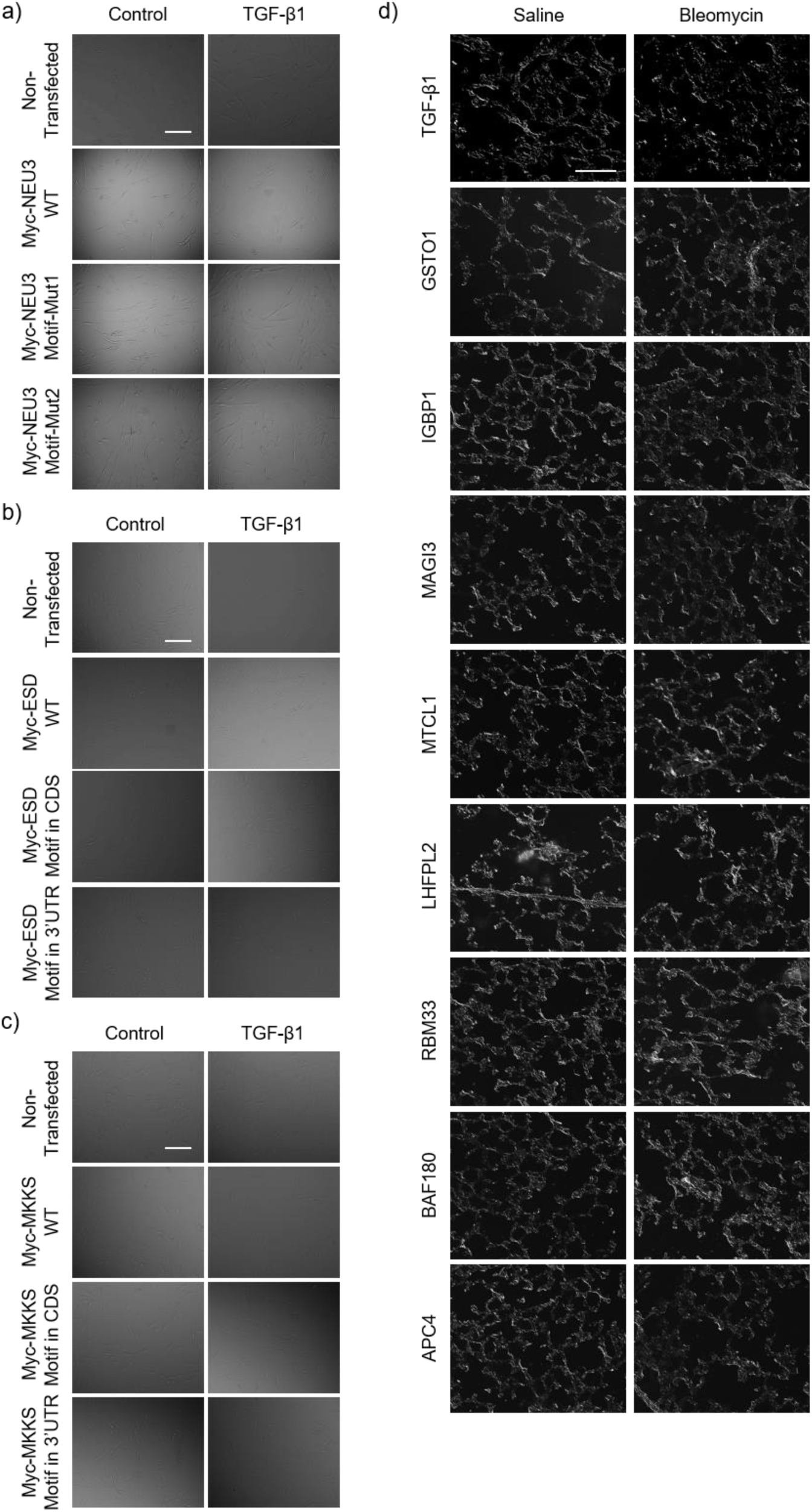
Corresponding DIC images of Figures 2a-c and 4e.

