## Supplemental file 7 for "Inhibiting a mRNA motif binding protein that mediates TGF-β1 upregulation of translation attenuates pulmonary fibrosis in mice"

1. Plasmid map of Myc-NEU3-WT:

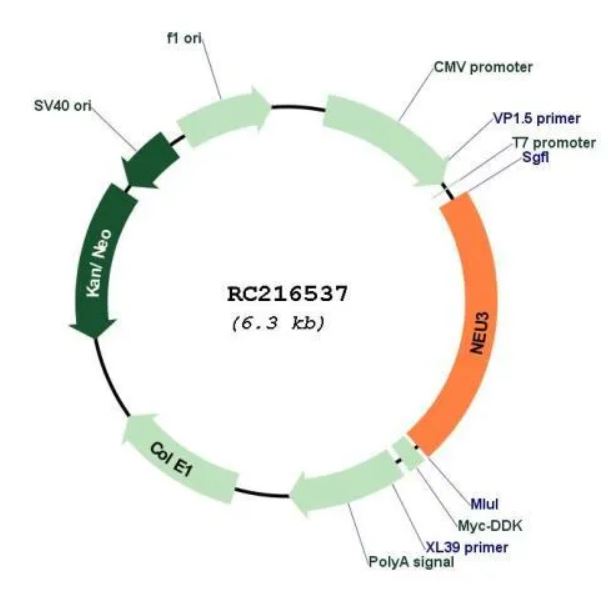

2. Nucleotide Sequences of Myc-NEU3-WT:

>RC216537 representing NM_006656

Red=Cloning site Blue=ORF Green=Tags(s)

TTTTGTAATACGACTCACTATAGGGCGGCCGGGAATTCGTCGACTGGATCCGGTACCGAGGAGATCTGCCGCCGCGATCGCCATGAGACCTGCGGACCTGCCCCCGCGCCCCATGGAAGAATCCCCGGCGTCCAGCTCTGCCCCGACAGAGACGGAGGAGCCGGGGTCCAGTGCAGAGGTCATGGAAGAAGTGACAACATGCTCCTTCAACAGCCCTCTGTTCCGGCAGGAAGATGACAGAGGGATTACCTACCGGATCCCAGCCCTGCTCTACATACCCCCCACCCACACCTTCCTGGCCTTTGCAGAGAAGCGTTCTACGAGGAGAGATGAGGATGCTCTCCACCTGGTGCTGAGGCGAGGGTTGAGGATTGGGCAGTTGGTACAGTGGGGGCCCCTGAAGCCACTGATGGAAGCCACACTACCGGGGCATCGGACCATGAACCCCTGTCCTGTATGGGAGCAGAAGAGTGGTTGTGTGTTCCTGTTCTTCATCTGTGTGCGGGGCCATGTCACAGAGCGTCAACAGATTGTGTCAGGCAGGAATGCTGCCCGCCTTTGCTTCATCTACAGTCAGGATGCTGGATGTTCATGGAGTGAGGTGAGGGACTTGACTGAGGAGGTCATTGGCTCAGAGCTGAAGCACTGGGCCACATTTGCTGTGGGCCCAGGTCATGGCATCCAGCTGCAGTCAGGGAGACTGGTCATCCCTGCGTATACCTACTACATCCCTTCCTGGTTCTTTTGCTTCCAGCTACCATGTAAAACCAGGCCTCATTCTCTGATGATCTACAGTGATGACCTAGGGGTCACATGGCACCATGGTAGACTCATTAGGCCCATGGTTACAGTAGAATGTGAAGTGGCAGAGGTGACTGGGAGGGCTGGCCACCCTGTGCTATATTGCAGTGCCCGGACACCAAACAGGTGCCGGGCAGAGGCGCTCAGCACTGACCATGGTGAAGGCTTTCAGAGACTGGCCCTGAGTCGACAGCTCTGTGAGCCCCCACATGGTTGCCAAGGGAGTGTGGTAAGTTTCCGGCCCCTGGAGATCCCACATAGGTGCCAGGACTCTAGCAGCAAAGATGCACCCACCATTCAGCAGAGCTCTCCAGGCAGTTCACTGAGGCTGGAGGAGGAAGCTGGAACACCGTCAGAATCATGGCTCTTGTACTCACACCCAACCAGTAGGAAACAGAGGGTTGACCTAGGTATCTATCTCAACCAGACCCCCTTGGAGGCTGCCTGCTGGTCCCGCCCCTGGATCTTGCACTGTGGGCCCTGTGGCTACTCTGATCTGGCTGCTCTGGAGGAGGAGGGCTTGTTTGGGTGTTTGTTTGAATGTGGGACCAAGCAAGAGTGTGAGCAGATTGCCTTCCGCCTGTTTACACACCGGGAGATCCTGAGTCACCTGCAGGGGGACTGCACCAGCCCTGGTAGGAACCCAAGCCAATTCAAAAGCAATACGCGTACGCGGCCGCTC**GAGCAGAAACTCATCTCAGAAGAGGATCTG**GCAGCAAATGATATCCTG**GATTACAAGGATGACGACGATAAG**GTTTAA

The two Group 4 motifs in NEU3 coding sequence are labeled with pink background.

3. Nucleotide Sequences of Myc-NEU3-Mutant1:

TTTTGTAATACGACTCACTATAGGGCGGCCGGGAATTCGTCGACTGGATCCGGTACCGAGGAGATCTGCCGCCGCGATCGCCATGAGACCTGCGGACCTGCCCCCGCGCCCCATGGAAGAATCCCCGGCGTCCAGCTCTGCCCCGACAGAGACGGAGGAGCCGGGGTCCAGTGCAGAGGTCATGGAAGAAGTGACAACATGCTCCTTCAACAGCCCTCTGTTCCGGCAGGAAGATGACAGAGGGATTACCTACCGGATCCCAGCCCTGCTCTACATACCCCCCACCCACACCTTCCTGGCCTTTGCAGAGAAGCGTTCTACGAGGAGAGATGAGGATGCTCTCCACCTGGTGCTGAGGCGAGGGTTGAGGATTGGGCAGTTGGTACAGTGGGGGCCCCTTAAACCACTTATTGAATCCACACTACCGGGGCATCGGACCATGAACCCCTGTCCTGTATGGGAGCAGAAGAGTGGTTGTGTGTTCCTGTTCTTCATCTGTGTGCGGGGCCATGTCACAGAGCGTCAACAGATTGTGTCAGGCAGGAATGCTGCCCGCCTTTGCTTCATCTACAGTCAGGATGCTGGATGTTCATGGAGTGAGGTGAGGGACTTGACTGAGGAGGTCATTGGCTCAGAGCTGAAGCACTGGGCCACATTTGCTGTGGGCCCAGGTCATGGCATCCAGCTGCAGTCAGGGAGACTGGTCATCCCTGCGTATACCTACTACATCCCTTCCTGGTTCTTTTGCTTCCAGCTACCATGTAAAACCAGGCCTCATTCTCTGATGATCTACAGTGATGACCTAGGGGTCACATGGCACCATGGTAGACTCATTAGGCCCATGGTTACAGTAGAATGTGAAGTGGCAGAGGTGACTGGGAGGGCTGGCCACCCTGTGCTATATTGCAGTGCCCGGACACCAAACAGGTGCCGGGCAGAGGCGCTCAGCACTGACCATGGTGAAGGCTTTCAGAGACTGGCCCTGAGTCGACAGCTCTGTGAGCCCCCACATGGTTGCCAAGGGAGTGTGGTAAGTTTCCGGCCCCTGGAGATCCCACATAGGTGCCAGGACTCTAGCAGCAAAGATGCACCCACCATTCAGCAGAGCTCTCCAGGCAGTTCACTGAGGCTGGAGGAGGAAGCTGGAACACCGTCAGAATCATGGCTCTTGTACTCACACCCAACCAGTAGGAAACAGAGGGTTGACCTAGGTATCTATCTCAACCAGACCCCCTTGGAGGCTGCCTGCTGGTCCCGCCCCTGGATCTTGCACTGTGGGCCCTGTGGCTACTCTGATCTGGCTGCTCTGGAGGAGGAGGGCTTGTTTGGGTGTTTGTTTGAATGTGGGACCAAGCAAGAGTGTGAGCAGATTGCCTTCCGCCTGTTTACACACCGGGAGATCCTGAGTCACCTGCAGGGGGACTGCACCAGCCCTGGTAGGAACCCAAGCCAATTCAAAAGCAATACGCGTACGCGGCCGCTC**GAGCAGAAACTCATCTCAGAAGAGGATCTG**GCAGCAAATGATATCCTG**GATTACAAGGATGACGACGATAAG**GTTTAA

5 sites were mutated in the 1^st^ Group 4 motif in NEU3 with the following mutation primers:

Forward	GGGCCCCTTAAACCACTTATTGAATCCACACTACC

Reverse	GGTAGTGTGGATTCAATAAGTGGTTTAAGGGGCCC

4. Nucleotide Sequences of Myc-NEU3-Mutant2:

TTTTGTAATACGACTCACTATAGGGCGGCCGGGAATTCGTCGACTGGATCCGGTACCGAGGAGATCTGCCGCCGCGATCGCCATGAGACCTGCGGACCTGCCCCCGCGCCCCATGGAAGAATCCCCGGCGTCCAGCTCTGCCCCGACAGAGACGGAGGAGCCGGGGTCCAGTGCAGAGGTCATGGAAGAAGTGACAACATGCTCCTTCAACAGCCCTCTGTTCCGGCAGGAAGATGACAGAGGGATTACCTACCGGATCCCAGCCCTGCTCTACATACCCCCCACCCACACCTTCCTGGCCTTTGCAGAGAAGCGTTCTACGAGGAGAGATGAGGATGCTCTCCACCTGGTGCTGAGGCGAGGGTTGAGGATTGGGCAGTTGGTACAGTGGGGGCCCCTGAAGCCACTGATGGAAGCCACACTACCGGGGCATCGGACCATGAACCCCTGTCCTGTATGGGAGCAGAAGAGTGGTTGTGTGTTCCTGTTCTTCATCTGTGTGCGGGGCCATGTCACAGAGCGTCAACAGATTGTGTCAGGCAGGAATGCTGCCCGCCTTTGCTTCATCTACAGTCAGGATGCTGGATGTTCATGGAGTGAGGTGAGGGACTTGACTGAGGAGGTCATTGGCTCAGAGCTGAAGCACTGGGCCACATTTGCTGTGGGCCCAGGTCATGGCATCCAGCTGCAGTCAGGGAGACTGGTCATCCCTGCGTATACCTACTACATCCCTTCCTGGTTCTTTTGCTTCCAGCTACCATGTAAAACCAGGCCTCATTCTCTGATGATCTACAGTGATGACCTAGGGGTCACATGGCACCATGGTAGACTCATTAGGCCCATGGTTACAGTAGAATGTGAAGTGGCAGAGGTGACTGGGAGGGCTGGCCACCCTGTGCTATATTGCAGTGCCCGGACACCAAACAGGTGCCGGGCAGAGGCGCTCAGCACTGACCATGGTGAAGGCTTTCAGAGACTGGCCCTGAGTCGACAGCTCTGTGAGCCCCCACATGGTTGCCAAGGGAGTGTGGTAAGTTTCCGGCCCCTGGAGATCCCACATAGGTGCCAGGACTCTAGCAGCAAAGATGCACCCACCATTCAGCAGAGCTCTCCAGGCAGTTCACTTAGACTGGAAGATGAATCTGGAACACCGTCAGAATCATGGCTCTTGTACTCACACCCAACCAGTAGGAAACAGAGGGTTGACCTAGGTATCTATCTCAACCAGACCCCCTTGGAGGCTGCCTGCTGGTCCCGCCCCTGGATCTTGCACTGTGGGCCCTGTGGCTACTCTGATCTGGCTGCTCTGGAGGAGGAGGGCTTGTTTGGGTGTTTGTTTGAATGTGGGACCAAGCAAGAGTGTGAGCAGATTGCCTTCCGCCTGTTTACACACCGGGAGATCCTGAGTCACCTGCAGGGGGACTGCACCAGCCCTGGTAGGAACCCAAGCCAATTCAAAAGCAATACGCGTACGCGGCCGCTC**GAGCAGAAACTCATCTCAGAAGAGGATCTG**GCAGCAAATGATATCCTG**GATTACAAGGATGACGACGATAAG**GTTTAA

5 sites were mutated in the 2^nd^ Group 4 motif in NEU3 with the following mutation primers:

Forward	CAGTTCACTTAGACTGGAAGATGAATCTGGAACAC

Reverse	GTGTTCCAGATTCATCTTCCAGTCTAAGTGAACTG

5. qPCR primers for Myc-NEU3 (cover both Myc-Tag and NEU3 sequences)

Forward	CAGATTGCCTTCCGCCTGTT

Reverse	CTGCCAGATCCTCTTCTGAGAT
