## Supplemental file 8 for "Inhibiting a mRNA motif binding protein that mediates TGF-β1 upregulation of translation attenuates pulmonary fibrosis in mice"

1. Plasmid map of Myc-ESD-WT

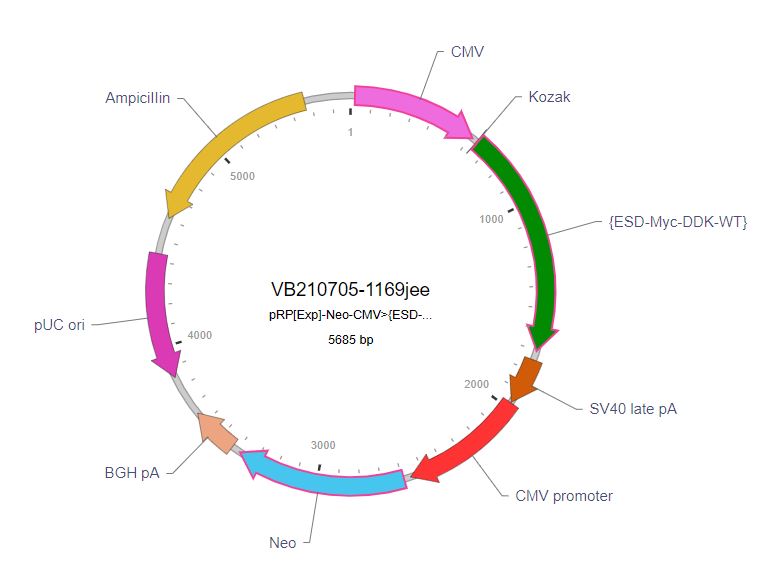

2. Nucleotide Sequences of Myc-ESD-WT:

aaagcgagagtgagtgggaccggaggggcggggcatcatatgggcggggctgaggcgaggccccggcggccatcttgagccccgccttttacttcggcccgcttcttctggtcactccgccaccgtagaatcgcctaccatttggtgcaagcaaaaagcaatcagcaattggacaggaaaagaATGGAGCAGAAACTCATCTCAGAAGAGGATCTGGATTACAAGGATGACGACGATAAGGCATTGAAGCAGATTTCCAGCAACAAGTGCTTTGGGGGATTGCAGAAAGTTTTTGAACATGACAGTGTTGAACTAAACTGCAAAATGAAATTTGCTGTCTACTTACCACCAAAGGCAGAAACAGGAAAGTGCCCTGCACTGTATTGGCTCTCAGGTTTAACTTGCACAGAGCAAAATTTTATATCAAAATCTGGTTATCATCAGTCTGCTTCAGAACATGGTCTTGTTGTCATTGCTCCAGATACCAGCCCTCGTGGCTGCAATATTAAAGGTGAAGATGAGAGCTGGGACTTTGGCACTGGTGCTGGATTTTATGTTGATGCCACTGAAGATCCTTGGAAAACCAACTACAGAATGTACTCTTATGTCACAGAGGAGCTTCCCCAACTCATAAATGCCAATTTTCCAGTGGATCCCCAAAGGATGTCTATTTTTGGCCACTCCATGGGAGGTCATGGAGCTCTGATCTGTGCTTTGAAAAATCCTGGAAAATACAAATCTGTGTCAGCATTTGCTCCAATTTGCAACCCTGTACTCTGTCCCTGGGGCAAAAAAGCCTTTAGTGGATATTTGGGAACAGATCAAAGTAAATGGAAGGCTTATGATGCTACCCACCTTGTGAAATCCTATCCAGGATCTCAGCTGGACATACTAATTGATCAAGGGAAAGATGACCAGTTTCTTTTAGATGGACAGTTACTCCCTGATAACTTCATAGCTGCCTGTACAGAAAAGAAAATCCCCGTTGTTTTTCGATTGCAAGAGGGTTATGATCATAGCTACTACTTCATTGCAACCTTTATTACTGACCACATCAGACATCATGCTAAATACCTGAATGCATGAaaaaactccaaataagagaatctcttcaggattataaaagttgtaaaatgcaactgtattgctgagcaaaaaaaaaaaaaattcaaaacattggattttatagtgctaaaagggctttattctatagttgaatcacctctgaataaagatataaaaccta

A Myc-Tag is “in-framely” inserted to the coding sequence of ESD.

3. Nucleotide Sequences of Myc-ESD-Motif in CDS:

aaagcgagagtgagtgggaccggaggggcggggcatcatatgggcggggctgaggcgaggccccggcggccatcttgagccccgccttttacttcggcccgcttcttctggtcactccgccaccgtagaatcgcctaccatttggtgcaagcaaaaagcaatcagcaattggacaggaaaagaATGGAGCAGAAACTCATCTCAGAAGAGGATCTGGATTACAAGGATGACGACGATAAGGCATTGAAGCAGATTTCCAGCAACAAGTGCTTTGGGGGATTGCAGAAAGTTTTTGAACATGACAGTGTTGAACTAAACTGCAAAATGAAATTTGCTGTCTACTTACCACCAAAGGCAGAAACAGGAAAGTGCCCTGCACTGTATTGGCTCTCAGGTTTAACTTGCACAGAGCAAAATTTTATATCAAAATCTGGTTATCATCAGTCTGCTTCAGAACATGGTCTTGTTGTCATTGCTCCAGATACCAGCCCTCGTGGCTGCAATATTAAAGGTGAAGATGAGAGCTGGGACTTTGGCACTGGTGCTGGATTTTATGTTGATGCCACTGAAGATCCTTGGAAAACCAACTACAGAATGTACTCTTATGTCACAGAGGAGCTTCCCCAACTCATAAATGCCAATTTTCCAGTGGATCCCCAAAGGATGTCTATTTTTGGAGGAGGAGGAGGAGGAGGCCACTCCATGGGAGGTCATGGAGCTCTGATCTGTGCTTTGAAAAATCCTGGAAAATACAAATCTGTGTCAGCATTTGCTCCAATTTGCAACCCTGTACTCTGTCCCTGGGGCAAAAAAGCCTTTAGTGGATATTTGGGAACAGATCAAAGTAAATGGAAGGCTTATGATGCTACCCACCTTGTGAAATCCTATCCAGGATCTCAGCTGGACATACTAATTGATCAAGGGAAAGATGACCAGTTTCTTTTAGATGGACAGTTACTCCCTGATAACTTCATAGCTGCCTGTACAGAAAAGAAAATCCCCGTTGTTTTTCGATTGCAAGAGGGTTATGATCATAGCTACTACTTCATTGCAACCTTTATTACTGACCACATCAGACATCATGCTAAATACCTGAATGCATGAaaaaactccaaataagagaatctcttcaggattataaaagttgtaaaatgcaactgtattgctgagcaaaaaaaaaaaaaattcaaaacattggattttatagtgctaaaagggctttattctatagttgaatcacctctgaataaagatataaaaccta

A Myc-Tag and a Group 4 motif are “in-framely” inserted to the coding sequence of ESD.

4. Nucleotide Sequences of Myc-ESD-Motif in 3’UTR:

aaagcgagagtgagtgggaccggaggggcggggcatcatatgggcggggctgaggcgaggccccggcggccatcttgagccccgccttttacttcggcccgcttcttctggtcactccgccaccgtagaatcgcctaccatttggtgcaagcaaaaagcaatcagcaattggacaggaaaagaATGGAGCAGAAACTCATCTCAGAAGAGGATCTGGATTACAAGGATGACGACGATAAGGCATTGAAGCAGATTTCCAGCAACAAGTGCTTTGGGGGATTGCAGAAAGTTTTTGAACATGACAGTGTTGAACTAAACTGCAAAATGAAATTTGCTGTCTACTTACCACCAAAGGCAGAAACAGGAAAGTGCCCTGCACTGTATTGGCTCTCAGGTTTAACTTGCACAGAGCAAAATTTTATATCAAAATCTGGTTATCATCAGTCTGCTTCAGAACATGGTCTTGTTGTCATTGCTCCAGATACCAGCCCTCGTGGCTGCAATATTAAAGGTGAAGATGAGAGCTGGGACTTTGGCACTGGTGCTGGATTTTATGTTGATGCCACTGAAGATCCTTGGAAAACCAACTACAGAATGTACTCTTATGTCACAGAGGAGCTTCCCCAACTCATAAATGCCAATTTTCCAGTGGATCCCCAAAGGATGTCTATTTTTGGCCACTCCATGGGAGGTCATGGAGCTCTGATCTGTGCTTTGAAAAATCCTGGAAAATACAAATCTGTGTCAGCATTTGCTCCAATTTGCAACCCTGTACTCTGTCCCTGGGGCAAAAAAGCCTTTAGTGGATATTTGGGAACAGATCAAAGTAAATGGAAGGCTTATGATGCTACCCACCTTGTGAAATCCTATCCAGGATCTCAGCTGGACATACTAATTGATCAAGGGAAAGATGACCAGTTTCTTTTAGATGGACAGTTACTCCCTGATAACTTCATAGCTGCCTGTACAGAAAAGAAAATCCCCGTTGTTTTTCGATTGCAAGAGGGTTATGATCATAGCTACTACTTCATTGCAACCTTTATTACTGACCACATCAGACATCATGCTAAATACCTGAATGCATGAaaaaactccaaataagagaatctcttcaggattataaaagttgtaaaatgcaactgtattgctgagcaaaaaaaaaaaaaattcaaaacattGGAGGAGGAGGAGGAGGAggattttatagtgctaaaagggctttattctatagttgaatcacctctgaataaagatataaaaccta

A Myc-Tag is “in-framely” inserted to the coding sequence of ESD. A Group 4 motif is “in-framely” inserted to 3’UTR of ESD.

5. qPCR primers for Myc-ESD (cover both Myc-Tag and ESD sequences)

Forward	GGATCTGGATTACAAGGATGAC

Reverse	CTGCCTTTGGTGGTAAGTAGAC
