## Supplemental file 9 for "Inhibiting a mRNA motif binding protein that mediates TGF-β1 upregulation of translation attenuates pulmonary fibrosis in mice"

1. Plasmid map of Myc-MKKS-WT

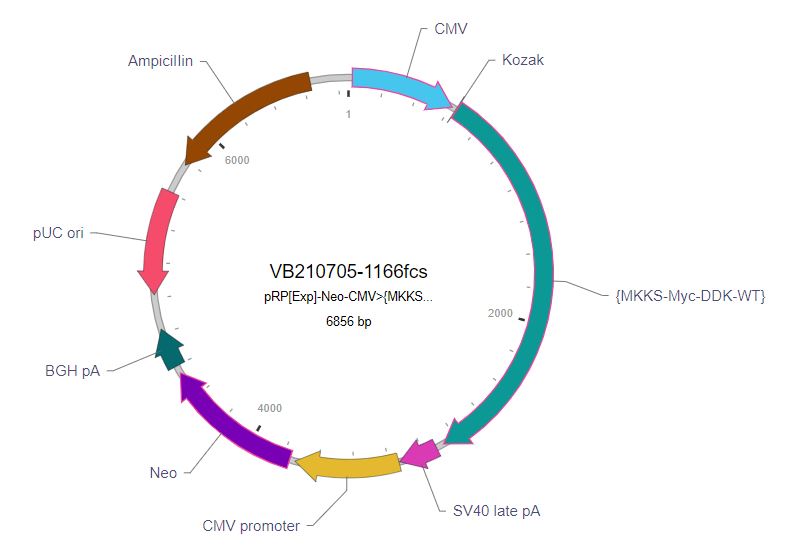

2. Nucleotide Sequences of Myc-MKKS-WT:

agagctgcgcgtgctccgtgccctcgcgcgacgcgaaggttgtcgggatccgcggcagcagcggctgcttgagatctgtttctggggcctctggcggtggcggcctggggcggcgcgacggctggtgcgcaggtacactgatgctgaagtactatgagccttcggaacttgtggagagactacaaagttttggttgttatggtccctttagttgggctcatacatttggggtggtacagaatcaaaagcagccctgttttccaaatacctaaaaacgacgacattcctgagcaagatagtctgggactttcaaatcttcagaagagccaaatccaggggaagtagcaggcttgcaatcttcaggtaaagaagcagctttgaatctgagcttcatatcgaaagaagagatgaaaaataccagttggattagaaagaactggcttcttgtagctgggatatctttcataggtgtccatcttggaacatactttttgcagaggtctgcaaagcagtctgtaaaatttcagtctcaaagcaaacaaaagagtattgaagagtgaagtaaaataaatatttggaattactaatttgtcattaaatcattctatgctgattagcttcataaacattgaactttttgattttatagccacaatgctgcatattcatactttaattcctaaagaataatttttaatgttaaaacgtgataatgcaataaatagaaaaatgtggtttacaaaataaaaacggtcttcactagttaccacctgaagtaagATGGAGCAGAAACTCATCTCAGAAGAGGATCTGGATTACAAGGATGACGACGATAAGTCTCGTTTGGAAGCTAAGAAGCCATCATTGTGTAAGAGTGAACCACTGACAACTGAGAGAGTCAGGACCACACTTTCTGTCTTGAAAAGAATTGTAACATCATGCTATGGCCCCTCAGGTAGGCTGAAGCAGCTGCACAATGGCTTTGGAGGTTACGTGTGTACAACCTCACAGTCCTCAGCTCTGCTCAGTCACCTTTTGGTCACACATCCCATTTTAAAGATCCTGACAGCCTCCATACAGAATCATGTGTCAAGCTTCAGTGATTGTGGCTTATTCACAGCTATTCTTTGCTGCAACCTGATTGAAAATGTTCAGAGATTAGGCTTGACACCCACCACTGTCATTAGATTAAATAAACATCTTTTGAGTCTTTGCATCAGTTATCTCAAGTCTGAGACCTGTGGTTGTCGAATCCCAGTGGACTTTAGTAGTACTCAGATCCTCCTTTGTTTGGTGCGTAGTATATTAACAAGTAAACCTGCCTGTATGCTCACCAGAAAGGAAACAGAGCATGTCAGTGCTTTGATCCTGAGAGCCTTTTTGCTTACAATTCCAGAAAATGCTGAAGGCCACATCATTTTAGGAAAGAGTTTAATTGTACCTTTAAAAGGTCAAAGAGTTATAGATTCCACTGTATTACCTGGGATACTCATTGAAATGTCAGAAGTTCAATTAATGAGGCTATTACCTATCAAAAAATCAACTGCCCTCAAGGTGGCACTCTTTTGTACAACTTTATCCGGAGACACTTCTGACACTGGAGAAGGAACTGTGGTGGTCAGTTATGGGGTTTCTCTTGAAAATGCAGTCTTGGACCAGCTGCTTAACCTAGGAAGGCAGCTAATCAGTGACCACGTAGATCTTGTCCTGTGCCAAAAAGTTATACATCCATCTTTGAAGCAGTTTCTCAATATGCATCGTATTATTGCCATAGACAGAATTGGAGTGACTCTGATGGAACCCCTGACTAAAATGACAGGAACACAGCCTATTGGATCCCTAGGCTCAATATGTCCTAATAGTTATGGAAGTGTGAAAGATGTGTGCACTGCAAAATTTGGCTCCAAACATTTTTTTCATCTTATTCCTAATGAAGCAACAATCTGCAGCTTGCTTCTCTGCAACAGAAATGACACTGCCTGGGATGAGCTGAAGCTCACGTGTCAGACGGCACTGCATGTCCTGCAGTTAACACTCAAGGAACCATGGGCTTTGTTGGGAGGTGGCTGTACTGAAACTCATTTGGCTGCATATATCAGACACAAGACTCACAACGACCCAGAAAGCATTCTCAAAGATGATGAATGTACTCAAACAGAACTTCAATTAATTGCTGAAGCATTTTGCAGTGCCCTAGAATCTGTTGTTGGCTCTTTAGAACATGATGGAGGTGAAATTCTCACTGACATGAAGTATGGACACCTTTGGTCAGTTCAGGCAGATTCTCCCTGTGTTGCTAACTGGCCAGATTTGCTTTCACAGTGTGGCTGTGGATTATACAATAGCCAGGAAGAACTCAACTGGTCTTTCTTAAGAAGCACACGTCGTCCATTTGTGCCACAAAGCTGCCTTCCACATGAAGCTGTGGGCTCAGCCAGCAACCTGACCTTGGACTGTTTGACTGCAAAGCTTAGTGGCCTACAGGTGGCTGTAGAGACAGCCAATTTGATTTTGGATCTTTCATATGTTATTGAAGATAAAAACTAAgagaatagcatgttcgtattacaagagaaacaaataaactagtctgttggcaattgagaaaaattgtgagtgtatttgttttctcccaaagccctgttctacatatttggacaaatgactcataaaattatagatacacttatttaggaaaaaaggtgattcgtgaatggaaatgccatgaaacaataaaaatatgaagcattatttatttaaaaatattatagttatcttagggattctatactggctgctgtacattgttctaaatttttgttatgttggcatcattttgagagcaaacaaataaaaaagactcctaatccatgctctagtttggatatacatattttagatattttccagttagcagtaattacatatgcttaaagtataaaactagatctcaaagtgtcacaaaacattcagtatatattgctcaatcaaaagaagtattcaaactgcacatt

A Myc-Tag is “in-framely” inserted to the coding sequence of MKKS.

3. Nucleotide Sequences of Myc-MKKS-Motif in CDS:

agagctgcgcgtgctccgtgccctcgcgcgacgcgaaggttgtcgggatccgcggcagcagcggctgcttgagatctgtttctggggcctctggcggtggcggcctggggcggcgcgacggctggtgcgcaggtacactgatgctgaagtactatgagccttcggaacttgtggagagactacaaagttttggttgttatggtccctttagttgggctcatacatttggggtggtacagaatcaaaagcagccctgttttccaaatacctaaaaacgacgacattcctgagcaagatagtctgggactttcaaatcttcagaagagccaaatccaggggaagtagcaggcttgcaatcttcaggtaaagaagcagctttgaatctgagcttcatatcgaaagaagagatgaaaaataccagttggattagaaagaactggcttcttgtagctgggatatctttcataggtgtccatcttggaacatactttttgcagaggtctgcaaagcagtctgtaaaatttcagtctcaaagcaaacaaaagagtattgaagagtgaagtaaaataaatatttggaattactaatttgtcattaaatcattctatgctgattagcttcataaacattgaactttttgattttatagccacaatgctgcatattcatactttaattcctaaagaataatttttaatgttaaaacgtgataatgcaataaatagaaaaatgtggtttacaaaataaaaacggtcttcactagttaccacctgaagtaagATGGAGCAGAAACTCATCTCAGAAGAGGATCTGGATTACAAGGATGACGACGATAAGTCTCGTTTGGAAGCTAAGAAGCCATCATTGTGTAAGAGTGAACCACTGACAACTGAGAGAGTCAGGACCACACTTTCTGTCTTGAAAAGAATTGTAACATCATGCTATGGCCCCTCAGGTAGGCTGAAGCAGCTGCACAATGGCTTTGGAGGTTACGTGTGTACAACCTCACAGTCCTCAGCTCTGCTCAGTCACCTTTTGGTCACACATCCCATTTTAAAGATCCTGACAGCCTCCATACAGAATCATGTGTCAAGCTTCAGTGATTGTGGAGGAGGAGGAGGAGGAGGCTTATTCACAGCTATTCTTTGCTGCAACCTGATTGAAAATGTTCAGAGATTAGGCTTGACACCCACCACTGTCATTAGATTAAATAAACATCTTTTGAGTCTTTGCATCAGTTATCTCAAGTCTGAGACCTGTGGTTGTCGAATCCCAGTGGACTTTAGTAGTACTCAGATCCTCCTTTGTTTGGTGCGTAGTATATTAACAAGTAAACCTGCCTGTATGCTCACCAGAAAGGAAACAGAGCATGTCAGTGCTTTGATCCTGAGAGCCTTTTTGCTTACAATTCCAGAAAATGCTGAAGGCCACATCATTTTAGGAAAGAGTTTAATTGTACCTTTAAAAGGTCAAAGAGTTATAGATTCCACTGTATTACCTGGGATACTCATTGAAATGTCAGAAGTTCAATTAATGAGGCTATTACCTATCAAAAAATCAACTGCCCTCAAGGTGGCACTCTTTTGTACAACTTTATCCGGAGACACTTCTGACACTGGAGAAGGAACTGTGGTGGTCAGTTATGGGGTTTCTCTTGAAAATGCAGTCTTGGACCAGCTGCTTAACCTAGGAAGGCAGCTAATCAGTGACCACGTAGATCTTGTCCTGTGCCAAAAAGTTATACATCCATCTTTGAAGCAGTTTCTCAATATGCATCGTATTATTGCCATAGACAGAATTGGAGTGACTCTGATGGAACCCCTGACTAAAATGACAGGAACACAGCCTATTGGATCCCTAGGCTCAATATGTCCTAATAGTTATGGAAGTGTGAAAGATGTGTGCACTGCAAAATTTGGCTCCAAACATTTTTTTCATCTTATTCCTAATGAAGCAACAATCTGCAGCTTGCTTCTCTGCAACAGAAATGACACTGCCTGGGATGAGCTGAAGCTCACGTGTCAGACGGCACTGCATGTCCTGCAGTTAACACTCAAGGAACCATGGGCTTTGTTGGGAGGTGGCTGTACTGAAACTCATTTGGCTGCATATATCAGACACAAGACTCACAACGACCCAGAAAGCATTCTCAAAGATGATGAATGTACTCAAACAGAACTTCAATTAATTGCTGAAGCATTTTGCAGTGCCCTAGAATCTGTTGTTGGCTCTTTAGAACATGATGGAGGTGAAATTCTCACTGACATGAAGTATGGACACCTTTGGTCAGTTCAGGCAGATTCTCCCTGTGTTGCTAACTGGCCAGATTTGCTTTCACAGTGTGGCTGTGGATTATACAATAGCCAGGAAGAACTCAACTGGTCTTTCTTAAGAAGCACACGTCGTCCATTTGTGCCACAAAGCTGCCTTCCACATGAAGCTGTGGGCTCAGCCAGCAACCTGACCTTGGACTGTTTGACTGCAAAGCTTAGTGGCCTACAGGTGGCTGTAGAGACAGCCAATTTGATTTTGGATCTTTCATATGTTATTGAAGATAAAAACTAAgagaatagcatgttcgtattacaagagaaacaaataaactagtctgttggcaattgagaaaaattgtgagtgtatttgttttctcccaaagccctgttctacatatttggacaaatgactcataaaattatagatacacttatttaggaaaaaaggtgattcgtgaatggaaatgccatgaaacaataaaaatatgaagcattatttatttaaaaatattatagttatcttagggattctatactggctgctgtacattgttctaaatttttgttatgttggcatcattttgagagcaaacaaataaaaaagactcctaatccatgctctagtttggatatacatattttagatattttccagttagcagtaattacatatgcttaaagtataaaactagatctcaaagtgtcacaaaacattcagtatatattgctcaatcaaaagaagtattcaaactgcacatt

A Myc-Tag and a Group 4 motif are “in-framely” inserted to the coding sequence of MKKS.

4. Nucleotide Sequences of Myc-ESD-Motif in 3’UTR:

agagctgcgcgtgctccgtgccctcgcgcgacgcgaaggttgtcgggatccgcggcagcagcggctgcttgagatctgtttctggggcctctggcggtggcggcctggggcggcgcgacggctggtgcgcaggtacactgatgctgaagtactatgagccttcggaacttgtggagagactacaaagttttggttgttatggtccctttagttgggctcatacatttggggtggtacagaatcaaaagcagccctgttttccaaatacctaaaaacgacgacattcctgagcaagatagtctgggactttcaaatcttcagaagagccaaatccaggggaagtagcaggcttgcaatcttcaggtaaagaagcagctttgaatctgagcttcatatcgaaagaagagatgaaaaataccagttggattagaaagaactggcttcttgtagctgggatatctttcataggtgtccatcttggaacatactttttgcagaggtctgcaaagcagtctgtaaaatttcagtctcaaagcaaacaaaagagtattgaagagtgaagtaaaataaatatttggaattactaatttgtcattaaatcattctatgctgattagcttcataaacattgaactttttgattttatagccacaatgctgcatattcatactttaattcctaaagaataatttttaatgttaaaacgtgataatgcaataaatagaaaaatgtggtttacaaaataaaaacggtcttcactagttaccacctgaagtaagATGGAGCAGAAACTCATCTCAGAAGAGGATCTGGATTACAAGGATGACGACGATAAGTCTCGTTTGGAAGCTAAGAAGCCATCATTGTGTAAGAGTGAACCACTGACAACTGAGAGAGTCAGGACCACACTTTCTGTCTTGAAAAGAATTGTAACATCATGCTATGGCCCCTCAGGTAGGCTGAAGCAGCTGCACAATGGCTTTGGAGGTTACGTGTGTACAACCTCACAGTCCTCAGCTCTGCTCAGTCACCTTTTGGTCACACATCCCATTTTAAAGATCCTGACAGCCTCCATACAGAATCATGTGTCAAGCTTCAGTGATTGTGGCTTATTCACAGCTATTCTTTGCTGCAACCTGATTGAAAATGTTCAGAGATTAGGCTTGACACCCACCACTGTCATTAGATTAAATAAACATCTTTTGAGTCTTTGCATCAGTTATCTCAAGTCTGAGACCTGTGGTTGTCGAATCCCAGTGGACTTTAGTAGTACTCAGATCCTCCTTTGTTTGGTGCGTAGTATATTAACAAGTAAACCTGCCTGTATGCTCACCAGAAAGGAAACAGAGCATGTCAGTGCTTTGATCCTGAGAGCCTTTTTGCTTACAATTCCAGAAAATGCTGAAGGCCACATCATTTTAGGAAAGAGTTTAATTGTACCTTTAAAAGGTCAAAGAGTTATAGATTCCACTGTATTACCTGGGATACTCATTGAAATGTCAGAAGTTCAATTAATGAGGCTATTACCTATCAAAAAATCAACTGCCCTCAAGGTGGCACTCTTTTGTACAACTTTATCCGGAGACACTTCTGACACTGGAGAAGGAACTGTGGTGGTCAGTTATGGGGTTTCTCTTGAAAATGCAGTCTTGGACCAGCTGCTTAACCTAGGAAGGCAGCTAATCAGTGACCACGTAGATCTTGTCCTGTGCCAAAAAGTTATACATCCATCTTTGAAGCAGTTTCTCAATATGCATCGTATTATTGCCATAGACAGAATTGGAGTGACTCTGATGGAACCCCTGACTAAAATGACAGGAACACAGCCTATTGGATCCCTAGGCTCAATATGTCCTAATAGTTATGGAAGTGTGAAAGATGTGTGCACTGCAAAATTTGGCTCCAAACATTTTTTTCATCTTATTCCTAATGAAGCAACAATCTGCAGCTTGCTTCTCTGCAACAGAAATGACACTGCCTGGGATGAGCTGAAGCTCACGTGTCAGACGGCACTGCATGTCCTGCAGTTAACACTCAAGGAACCATGGGCTTTGTTGGGAGGTGGCTGTACTGAAACTCATTTGGCTGCATATATCAGACACAAGACTCACAACGACCCAGAAAGCATTCTCAAAGATGATGAATGTACTCAAACAGAACTTCAATTAATTGCTGAAGCATTTTGCAGTGCCCTAGAATCTGTTGTTGGCTCTTTAGAACATGATGGAGGTGAAATTCTCACTGACATGAAGTATGGACACCTTTGGTCAGTTCAGGCAGATTCTCCCTGTGTTGCTAACTGGCCAGATTTGCTTTCACAGTGTGGCTGTGGATTATACAATAGCCAGGAAGAACTCAACTGGTCTTTCTTAAGAAGCACACGTCGTCCATTTGTGCCACAAAGCTGCCTTCCACATGAAGCTGTGGGCTCAGCCAGCAACCTGACCTTGGACTGTTTGACTGCAAAGCTTAGTGGCCTACAGGTGGCTGTAGAGACAGCCAATTTGATTTTGGATCTTTCATATGTTATTGAAGATAAAAACTAAgagaatagcatgttcgtattacaagagaaacaaataaactagtctgttggcaattgagaaaaattgtgagtgtatttgttttctcccaaagccctgttctacatatttGGAGGAGGAGGAGGAGGAggacaaatgactcataaaattatagatacacttatttaggaaaaaaggtgattcgtgaatggaaatgccatgaaacaataaaaatatgaagcattatttatttaaaaatattatagttatcttagggattctatactggctgctgtacattgttctaaatttttgttatgttggcatcattttgagagcaaacaaataaaaaagactcctaatccatgctctagtttggatatacatattttagatattttccagttagcagtaattacatatgcttaaagtataaaactagatctcaaagtgtcacaaaacattcagtatatattgctcaatcaaaagaagtattcaaactgcacatt

A Myc-Tag is “in-framely” inserted to the coding sequence of MKKS. A Group 4 motif is “in-framely” inserted to 3’UTR of MKKS.

5. qPCR primers for Myc-MKKS (cover both Myc-Tag and MKKS sequences)

Forward	CATCTCAGAAGAGGATCTGG

Reverse	CAAGACAGAAAGTGTGGTCC
