## Supplementary figures and images for "Inhibiting a mRNA motif binding protein that mediates TGF-β1 upregulation of translation attenuates pulmonary fibrosis in mice"

### Supplemental file 10

1. Biotin-Group 4 Motif sequence:

/5Biosg/ GGAGGAGGAGGAGGAGGAGG

2. 5’ Biotin information:

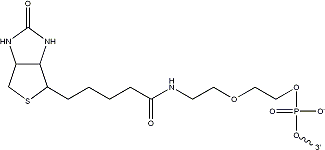

MW: 393.4
